# Interneuron migration defects during corticogenesis contribute to *Dyrk1a* haploinsufficiency syndrome pathogenesis via actomyosin dynamics deregulations

**DOI:** 10.1101/2023.11.09.566424

**Authors:** Maria Victoria Hinckelmann, Aline Dubos, Victorine Artot, Gabrielle Rudolf, Thu Lan Nguyen, Peggy Tilly, Valérie Nalesso, Maria del Mar Muniz Moreno, Marie-Christine Birling, Juliette D. Godin, Véronique Brault, Yann Herault

## Abstract

Interneuron development is a crucial step of brain corticogenesis. When affected it often leads to brain dysfunctions, such as epilepsy, intellectual disabilities and autism spectrum disorder. Such defects are observed in the *DYRK1A*-haploinsufficiency syndrome, caused by mutations of *DYRK1A*, and commonly associated to cortical excitatory/inhibitory imbalance. However, how this imbalance is established in this syndrome remains elusive. Here, using mouse models and live imaging, we show that *Dyrk1a* specifically regulates the development of the cortical GABAergic system. Unlike projection excitatory neurons, we demonstrate that interneuron tangential migration relies on Dyrk1a dosage and kinase activity through a mechanism involving actomyosin cytoskeleton remodeling. Interestingly, we further demonstrate that mice with heterozygous inactivation of *Dyrk1a* in interneurons show behavioral defects and epileptic activity, recapitulating phenotypes observed in human patients. Altogether, these data highlight the critical role of *Dyrk1a* in the development of the GABAergic system and the pathophysiology of *DYRK1A*-haploinsufficiency syndrome.

## INTRODUCTION

Interneurons (INs) represent a small population of neurons in the brain whose role is key to assure normal adult brain functioning. Several genetic diseases have been described with consequences on the development of INs causing a shift in the excitatory/inhibitory balance in the adult brain. This balance is known to regulate neurophysiological and cognitive processes ^1,2^, and its deregulation leads to pathological conditions such as ID ^3–5^, ASD ^6–8^ and epilepsy ^9^. As such, cortical INs play a crucial role in regulating the excitation-inhibition balance and in fine-tuning the functioning of cortical networks. The establishment of this functional balance in the brain begins during embryogenesis, at the time when excitatory and inhibitory neurons are produced.

Over the last fifteen years, different variants affecting the function of *DYRK1A* (dual-specificity tyrosine phosphorylation regulated kinase 1A) have been identified among rare individuals with a syndromic form of intellectual disability (ID). These mutations - which include point mutations, insertions or deletions (indels), chromosomal rearrangements, and microindels - affect one *DYRK1A* allele, and result in a distinctive syndrome known as *DYRK1A*-haploinsufficiency syndrome (DHS) or Mental Retardation Dominant 7 (MRD7, OMIM: #614104). DHS is a rare neurodevelopmental disorder characterized by moderate to severe ID, primary microcephaly, speech impairments and motor delay, distinctive facial dysmorphism, and is frequently associated with conditions such as autism spectrum disorder (ASD) and epilepsy ^10–18^.

*DYRK1A* encodes a dual-specificity tyrosine (Y) phosphorylation-regulated kinase, with both nuclear and cytoplasmic substrates and interactors ^19,20^. Through these functional interactions DYK1A regulates a number of biological processes such as transcription, cell cycle, cellular differentiation, vesicle trafficking and survival (reviewed in ^21^). DYRK1A is a constitutively active protein that requires the autophosphorylation of a critical tyrosine residue (Y321) located in the kinase domain of the protein to achieve its full catalytic activity ^22–24^.

Since its discovery with *minibrain* (*Mnb*) *Drosophila* mutant ^25^ and its homolog on human chromosome 21 ^26^, *DYRK1A* has been extensively studied and implicated in the correct development and functioning of the brain. *DYRK1A* is considered as a main driver of the neurodevelopmental alterations that lead to the cognitive impairments observed in Down Syndrome (DS) ^20,27^. Indeed, *Dyrk1a* is expressed in fetal and adult brain ^28–30^, supporting a role of *DYRK1A* as a regulator of brain growth and function. It was shown to control different processes such as neurogenesis, neuronal proliferation, differentiation, synaptic transmission and death ^20,21^. For instance, mutations in *Mnb* result in the reduction of the brain size of adult flies due to defective neuroblast proliferation and neurogenesis ^25^. Similar phenotypes in brain size, cellularity and neurogenesis were also observed in heterozygous knock-out mouse models, which were associated with behavioral alterations including seizures, cognition and sociability impairments ^31–35^, resembling the human DHS pathology.

Interestingly, alterations in *Dyrk1a* gene dosage disrupt the proportion of GABAergic inhibitory and glutamatergic excitatory systems in adult mouse cortices ^36^. This leads to opposite excitatory/inhibitory (E/I) imbalance in the brain: towards inhibition in the case of *Dyrk1a* overexpression, and towards excitation in *Dyrk1a* haploinsufficiency ^31,33,37,38^. Although, it is well known that *Dyrk1a* regulates the development of the cortical excitatory system ^36,39–41^, its physiological function in INs development remains relatively unexplored. Here, we analyzed the cell-autonomous contribution of the GABAergic system in the pathophysiology of DHS. We report that *Dyrk1a* haploinsufficient INs fail to efficiently invade the developing cortex due to defects in tangential migration, which can be attributed to reduced kinase activity of DYRK1A. These defects are associated with altered actomyosin dynamics at the rear of the nucleus during nucleokinesis. Finally, we show that the specific inactivation of *Dyrk1a* in the GABAergic system leads to cognitive and social deficits associated with an increased susceptibility to epilepsy in adult mice, mirroring human DHS phenotypes.

## RESULTS

### *Dyrk1a* haploinsufficiency affects interneurons distribution in the developing cortex

To analyze the cell autonomous contribution of *Dyrk1a* in the development of cortical INs we crossed the Tg(dlx5a-cre)1 ^Mekk^ (IMSR_JAX:008199) mice -named hereafter *DlxCre* ^42^-which express the CRE recombinase in postmitotic GABAergic INs, with a conditional knock-out mouse for *Dyrk1a*, *Dyrk1a ^lox/+^* ^27^. In this way, we deleted one gene copy of *Dyrk1a* specifically in postmitotic INs (*Dyrk1a^DlxCre/+^*). To visualize GABAergic INs, we further bred these mice with the transgenic GAD65-EGFP model expressing the green fluorescent protein GFP under the control of the INs specific promoter of the GAD65 gene ^43^ (Fig. 1a).

**Figure 1:**
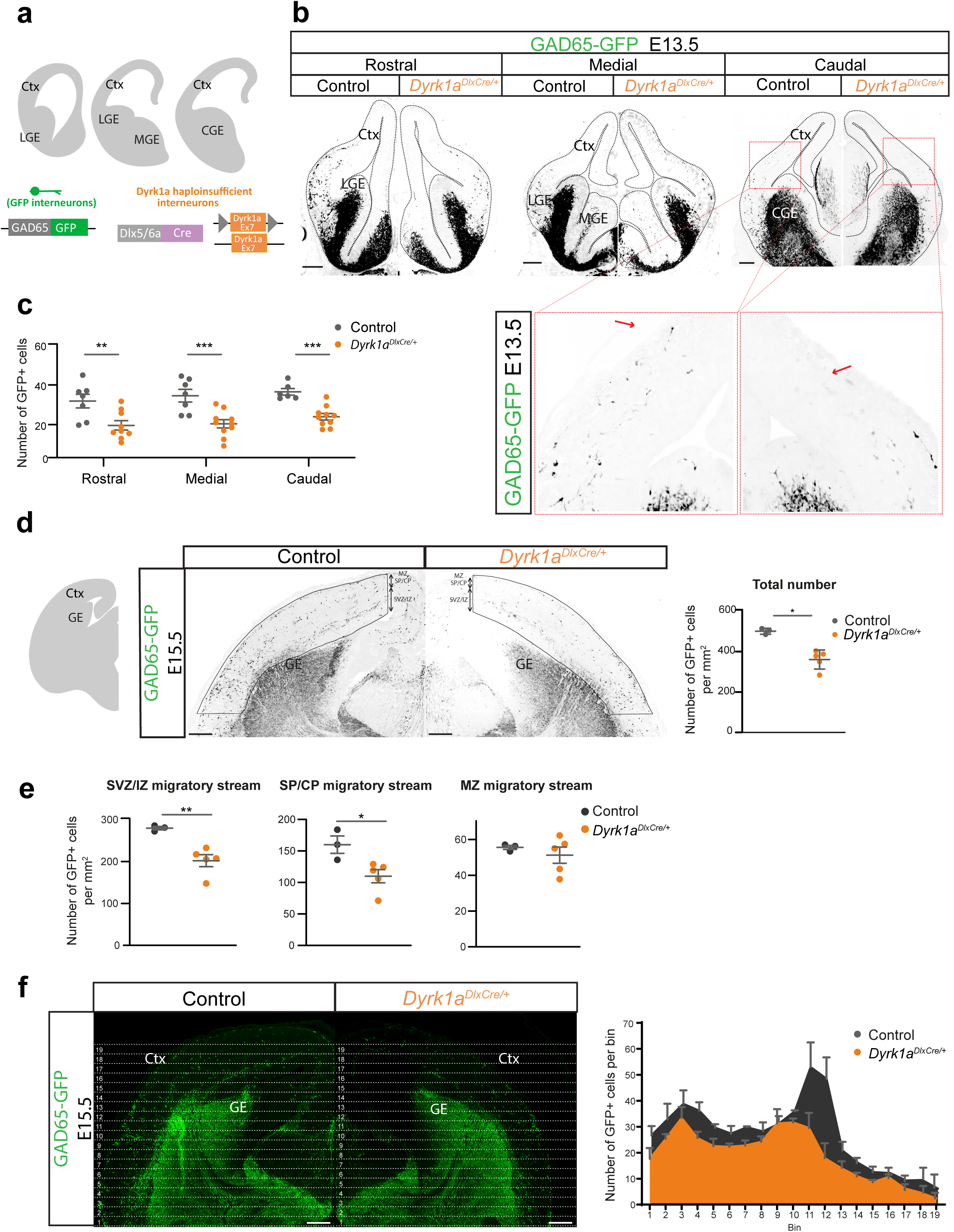
*Dyrk1a* haploinsufficiency affects interneurons distribution in the developing cortex. **a** Schematic representation of the three different E13.5 rostro-caudal sections of the brain analyzed. *Dyrk1a* conditional allele was used to selectively remove one copy of the gene in postmitotic INs. INs were visualized thanks to the expression of the GFP under the GAD65 promoter. **b** Representative E13.5 brain sections of control and *Dyrk1a^DlxCre/+^* embryos. GFP-expressing INs (black) migrate from the GEs into the Ctx. The brain is contoured with black dashed lines. Higher magnification of the region inside the red dashed line. Red arrows show the migration front. Scale bar, 200µm. **c** Quantification of the number of GFP+ INs in control and *Dyrk1a^DlxCre/+^* brains entering the developing cortex in three different rostro-caudal levels. n=8 embryos (control), n=10 embryos (*Dyrk1a^DlxCre/+^*) from at least three different mothers, two-way ANOVA, **p < 0.01, ***p < 0.001. **d** Schematic representation of the E15.5 brain sections analyzed (left panel). Representative coronal sections of E15.5 control and *Dyrk1a^DlxCre/+^*brains, showing GAD65-EGFP INs (black). Scale bar, 200µm. Quantifications show the number of GFP+ INs per mm ^2^ in E15.5 developing cortices. n=3 embryos (control), n=5 embryos (*Dyrk1a^DlxCre/+^*) from at least three different mothers, unpaired t-test, *p < 0.05. **e** Distribution of INs among the different migratory streams in the E15.5 developing cortex, shown as the number of GFP+ cells per mm ^2^ located either in the SVZ/VS, SP/CP or MZ migratory stream. n=3 embryos (control), n=5 embryos (*Dyrk1a^DlxCre/+^*) from at least three different mothers, unpaired t-test, *p < 0.05, **p < 0.01. **f** Representative brain sections of E15.5 embryonic brains (left panel). Scale bar, 200 µm. Sections were divided in 19 dorso-ventral bins of the same size. The number of GFP+ INs located in each bin was quantified (right panel). n=3 embryos (control), n=5 embryos (*Dyrk1a^DlxCre/+^*) from at least three different mothers, unpaired t-test, *p < 0.05. Data is represented as dot plots with mean ± s.e.m. CP, cortical plate; Ctx, cortex; LGE, lateral ganglionic eminence; MGE, medial ganglionic eminence; CGE, caudal ganglionic eminence; SVZ, subventricular zone; VZ, ventricular zone; SP, subplate; MZ, marginal zone.

Cortical INs are produced in the ganglionic eminences (GEs) where their progenitors reside, and at around 13 days post-coitum (E13.5) they begin to migrate out of these structures towards the developing cortex. *Dyrk1a* is expressed in the GEs during neurogenesis ^28^, and its protein is present in the soma and processes of migrating INs (Supplementary Fig. 1a). At this stage, *Dyrk1a* haploinsufficiency in INs leads to defects in their ability to exit the GEs. Less GFP-expressing INs crossed the cortico-striatal boundary (CSB) and entered the cortex, and the migration front of invading INs was delayed (Fig. 1b) . This reduction affected similarly all three studied rostro-caudal regions of the developing brain (Fig. 1a, b, c). Later in development, at E15.5, the number of INs that reached the cortex in *Dyrk1a^DlxCre/+^* sections was significantly reduced compared to controls (Fig. 1d), showing a persistent defect in the number of INs reaching the cortex. These alterations were observed in the deep migratory streams -intermediate zone/subventricular zone (IZ/SVZ), and the subplate and cortical plate (SP/CP)-, whereas the INs migrating through the superficial marginal zone (MZ) migratory stream were not affected (Fig. 1e). Then, we assessed whether this reduction in the number of INs populating the cerebral cortex was due to increased cell death or non-cell autonomous effects on progenitors proliferation. We observed no difference in the number of dividing progenitors in the different GEs of E12.5 and E15.5 embryos (Supplementary Fig. 1b, c), and no evident increase in apoptosis at these embryonic stages in *Dyrk1a^DlxCre/+^* brains (Supplementary Fig. 1d, e). This suggests that *Dyrk1a* haploinsufficiency does not affect the survival of newly generated nor migrating INs, and as expected excludes non-cell-specific effects on IN progenitors. We next tested cell autonomous effect of *Dyrk1a* haploinsufficiency on migration. We analyzed the distribution of GFP+ cells in coronal sections of E15.5 brains by dividing the cortex dorso-ventrally in 19 equal bins and scoring the number of GFP+ cells in each bin. As such, the low numbered bins represent INs that just exited the GE whereas those that have advanced in their migration will be located in the high numbered bins. The distribution profile obtained clearly shows a delay in the migration front. Indeed, besides the overall reduction of INs within *Dyrk1a^DlxCre/+^*cortices, there is a shift in the peak of GFP+ cells towards more delayed bins (or lower number bins) (Fig. 1f) in haploinsufficient INs. Together, this data suggests that the reduced number of INs in *Dyrk1a* haploinsufficient cortices likely stems from a migration defect of INs during development.

As *Dyrk1a* post-mitotically controls the distribution of INs in the cortex, we wondered whether *Dyrk1a* could also regulate the distribution of pyramidal neurons. To study this, we performed *in utero* electroporation to remove one copy of *Dyrk1a* in postmitotic pyramidal neurons. E14.5 mouse cortices from *Dyrk1a ^lox/+^* embryos were electroporated with a construct expressing the CRE-recombinase and the GFP under the neuron-specific promoter NeuroD, and four days later (E18.5) the distribution of GFP+ electroporated cells was analyzed (Supplementary Fig. 2a). Contrary to what we observed in GABAergic INs, we found no defects in the laminar organization of *Dyrk1a* haploinsufficient projecting neurons (Supplementary Fig. 2b, c). Eventually, to exclude any involvement of *Dyrk1a* in postmitotic pyramidal neuron migration, we performed the same analyses on *Dyrk1a ^lox/lox^* embryos and we did not observe any defect in postmitotic pyramidal neuron distribution in the complete absence of *Dyrk1a* (Supplementary Fig. 2b, d). Together, these observations indicate that *Dyrk1a* regulates postmitotic processes controlling the cell cycle exit or migration processes of GABAergic INs during corticogenesis in a cell-autonomous manner.

### *Dyrk1a* haploinsufficiency leads to abnormal localization of interneurons after birth

The development of the GABAergic system continues during the first weeks after birth. During this period the laminar allocation of tangentially migrating INs, subtype refinement and integration into brain circuits occur ^44,45^. So, we next wanted to understand whether and how the delay in the migration front during embryogenesis could affect the organization and development of the GABAergic system at early postnatal stages. To study this, we first quantified and analyzed INs distribution in the dorso-lateral cortex at different rostro-caudal levels of the brain (Fig. 2a). We observed that the decrease in GABAergic cells was still present 8 days after birth (P8), with a net reduction in the density of GAD65-GFP+ cells (Fig. 2 b, c). Next, to evaluate the intracortical dispersion of GAD65-GFP+ INs, we divided the cortical wall in 10 bins of the same size, and analyzed the distribution of GAD65-GFP+ INs within these bins. We observed that, although the density of GFP+ cells is decreased (Fig. 2c), the relative distribution of GFP+ cells within the cortical wall was similar between control and *Dyrk1a* haploinsufficient INs (Fig. 2b, d), suggesting that the decrease of GAD65-GFP+ INs is not specific of one bin. INs diversity depends on both the origin and timing of the production of these cells ^44,46,47^. Then, to assess potential differences in INs subtypes due to *Dyrk1a* inactivation, we studied the distribution and laminar organization of INs subpopulations. In control brains, as expected, cell density was specific to the different IN subtypes, ranging around 150 cells/mm ^2^ for GAD65-GFP+ and calbindin (CB+) cells and 75 cells/mm^2^ for somatostatin (SST+) INs (Fig. 2b-c, e-h) . Additionally, the distribution pattern within the cortical wall also displayed distinctive features corresponding to each IN marker (Fig. 2b, d, Supplementary Fig. 3a, b). However, as with GAD65-GFP+ INs, *Dyrk1a* downregulation also affected CB+ and SST+ INs. In *Dyrk1a^DlxCre/+^*cortices, we found a reduction in the density of both CB+ and SST+ INs subtypes (Fig. 2e, f, g, h) . Still (Fig. 2b, d), the distribution within the cortical wall of both INs markers was not differentially affected (Supplementary Fig. 3a, b).

**Figure 2:**
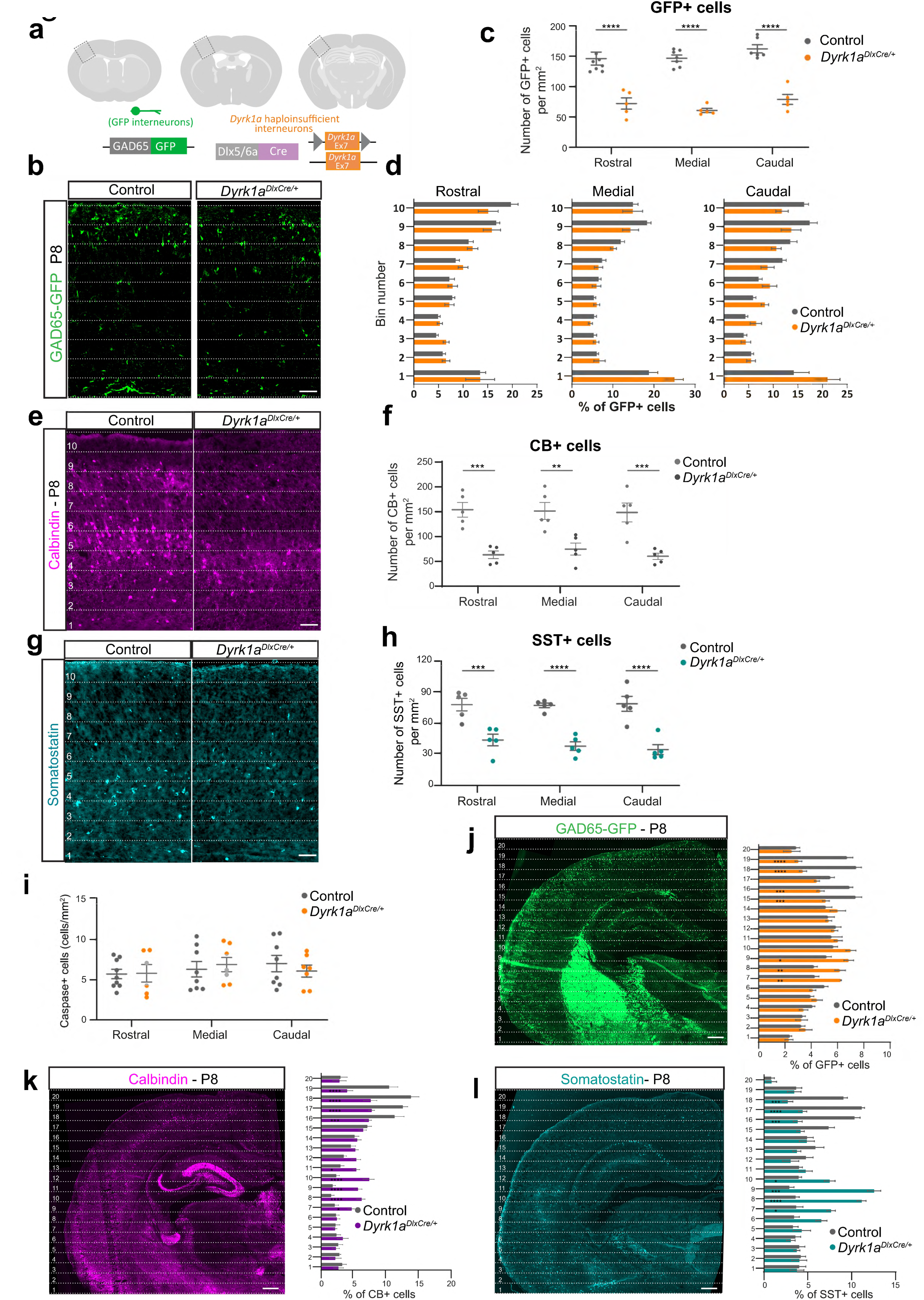
*Dyrk1a* haploinsufficiency leads to abnormal localization of interneurons after birth. **a** Schematic representation of the three different P8 rostro-caudal brain sections analyzed. The quantification area is shown within a dotted-line. *Dyrk1a* conditional allele was used to selectively remove one copy of the gene in postmitotic INs. INs were visualized thanks to the expression of the GFP under the GAD65 promoter. **b** Representative images of the dorso-lateral cortex of postnatal day 8 control and *Dyrk1a^DlxCre/+^* pups. GAD65-GFP INs are shown in green. The cortex was divided in 10 equal bins. Scale bar, 100µm. **c** Number of GFP+ INs per mm ^2^ in control and *Dyrk1a^DlxCre/+^* dorso-lateral cortices, at three different rostro-caudal positions. n=7 brains (control), n=5 brains (*Dyrk1a^DlxCre/+^*) from at least three different mothers, unpaired t-test, ****p < 0.0001. **d** Histograms represent the percentage of GFP-positive INs within each bin at three different rostro-caudal positions in the cortex. n=7 embryos (control), n=5 embryos (*Dyrk1a^DlxCre/+^*) from at least three different mothers, two-way ANOVA. **e** Representative images of the dorso-lateral cortex of postnatal day 8 control and *Dyrk1a^DlxCre/+^* pups. Sections were stained against calbindin (magenta). The cortex was divided in 10 equal bins. Scale bar, 100µm. **f** Graph represents the density of calbindin-positive INs per mm ^2^. n=5 embryos (control), n=4 embryos (*Dyrk1a^DlxCre/+^*) from at least three different mothers, unpaired t-test, **p < 0.01, ****p < 0.0001. **g** Representative images of the dorso-lateral cortex of postnatal day 8 control and *Dyrk1a^DlxCre/+^* pups. Sections were stained against somatostatin (cyan). The cortex was divided in 10 equal bins. Scale bar, 100µm. **h** Graph represents the density of somatostatin-positive INs within the dorso-lateral cotex. n=6 embryos (control), n=5 embryos (*Dyrk1a^DlxCre/+^*) from at least three different mothers, unpaired t-test, ***p < 0.001, ****p < 0.0001. **i** Quantification of caspase positive INs within the cortex in control and *Dyrk1a^DlxCre/+^* brains. n=8 embryos (control), n=5 embryos (*Dyrk1a^DlxCre/+^*) from at least three different mothers, unpaired t-test. **j** Coronal section of postnatal day 8 pups were dorso-ventrally divided in 20 equal bins. GAD65-GFP INs are shown in green (left panel). Scale bar, 200µm. The number of GFP-positive INs in the cortex within each bin was quantified in control and *Dyrk1a^DlxCre/+^* cortices (right panel). **k** Coronal section of postnatal day 8 pups were dorso-ventrally divided in 20 equal bins. Sections were stained against calbindin (magenta) (left panel). Scale bar, 200µm. The number of calbindin-positive INs within each bin was quantified in control and *Dyrk1a^DlxCre/+^*cortices (right panel). **l** Coronal section of postnatal day 8 pups were dorso-ventrally divided in 20 equal bins. Sections were stained for somatostatin (cyan) (left panel). Scale bar, 200µm. The number of somatostatin-positive INs within each bin was quantified in control and *Dyrk1a^DlxCre/+^* cortices (right panel). Data is represented dot plots with mean ± s.e.m, or histograms with mean ± s.e.m. P8, postnatal day 8; CB, calbindin; SST, somatostatin.

During corticogenesis there is a physiological overproduction of INs. Therefore, between the first and second week after birth, excess INs are normally eliminated through programmed cell death ^48–50^. However, though *Dyrk1a* has been involved in the regulation of apoptosis ^21^, cell death alterations did not account for the decrease of INs in the somatosensory cortex in *Dyrk1a^DlxCre/+^* brains at P8 (Fig. 2i). On the other hand, this depletion of GFP+ cells in the dorsal cortex was associated with an accumulation of INs in regions of the cortex located ventrally in the brain (Fig. 2j). In *Dyrk1a^DlxCre/+^* cortices, there is a significant increase of GAD65-GFP+ IN in lower number bins with a depletion of these cells in higher number bins, reminiscent of the delay in the migration front described at E15.5 (Fig. 1f). This reorganization, was also observed in different INs subtypes. Both CB+ (Fig. 2k) and SST+ (Fig. 2l) INs are redistributed with an increased localization in ventral areas of the cortex, with differences that correspond to their specific distribution within the cortex.

Altogether, this indicates that the delay in migration is not a transient phenomenon, and suggests that INs that were left behind during tangential migration were unable to catch up and reach their proper localization in the cortex.

### *Dyrk1a* dosage controls interneurons migration dynamics

To examine the consequences of *Dyrk1a* haploinsufficiency in the migration of GABAergic INs, we studied tangential migration using time-lapse video-microscopy. This allows us to follow the dynamics in morphology and locomotion of migrating INs exiting the GEs. We cultured E13.5 control and *Dyrk1a^DlxCre/+^* caudal GEs (CGE)-derived explants (from where most of the cortical GAD65-GFP+ INs are produced) on top of wild-type cortical feeders, and imaged migrating INs one day later during 5 hours (Fig. 3a). Acquisitions showed that in both conditions INs were able to migrate out of the explants and pursue active migration (Fig. 3b, c). However, haploinsufficient INs moved slower. We tracked single migrating cells, and observed that their displacement velocity was significantly lower (Fig. 3c, d), and that the duration of pauses was concomitantly increased (Fig. 3e). INs typically use a saltatory mode of migration that consists of cycles of nuclear displacements (nucleokinesis) and pauses ^51^ (Fig. 3f). As this movement is essential for locomotion, we analyzed in detail the motion of these nuclei during INs migration. We found that *Dyrk1a* haploinsufficiency leads to a significant decrease in both the frequency and amplitude of nucleokinesis (Fig. 3g, h), which likely contributes to the net decrease in migration velocity of these cells. Cortical INs derive mostly from the CGE, MGE and the preoptic region. Each specific domain generates in a temporally-restricted manner different IN subtypes during corticogenesis ^44,46,47^. Therefore, to evaluate whether *Dyrk1a* governs INs dynamics, regardless of their origin, we crossed the conditional *Dyrk1a^lox/+^* allele with *Dlx-Cre-IRES-GFP* mice (Tg(mI56i-cre,EGFP)1 ^Kc^; IMSR JAX:023724) ^52^. This is a frequently used model that allows the concomitant expression of CRE-recombinase with the GFP in postmitotic INs derived in this case from the MGE (Supplementary Fig. 4a). Time-lapse experiments from MGE-derived explants demonstrate that the dynamics of migration of GABAergic INs originating from the MGE was similarly affected in *Dyrk1a^DlxCreIRESGFP/+^*embryos (Supplementary Fig. 4b-e).

**Figure 3:**
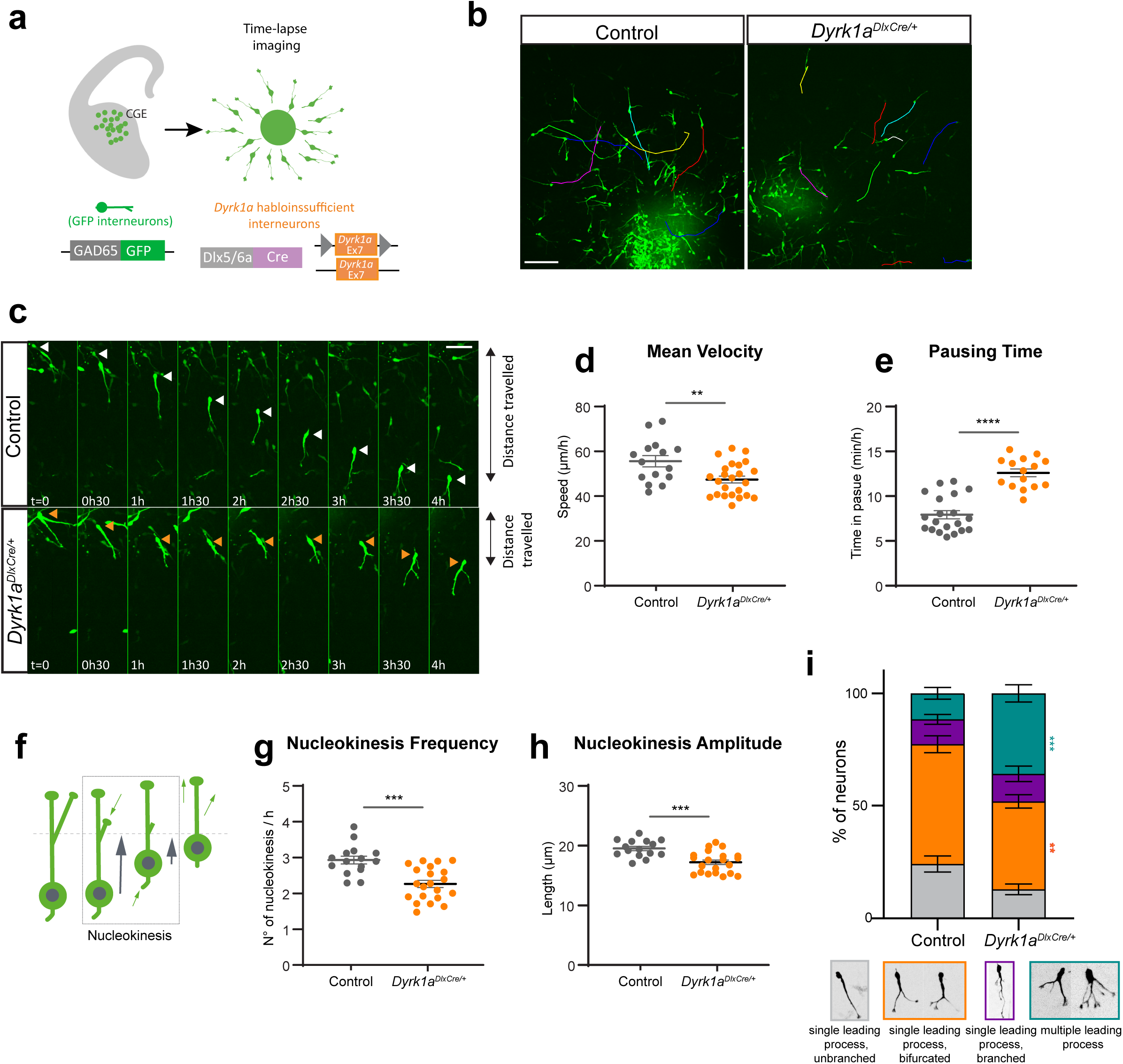
*Dyrk1a* dosage controls INs migration dynamics and morphology. **a** Caudal ganglionic eminences explants of E13.5 control and *Dyrk1a^DlxCre/+^* embryos were cultured on top of wild type cortical feeders. One day later GFP+ INs exiting the explants were imaged every 5min for 5 hours. **b** Migration trajectories (colored lines) of single GFP+ migrating INs recorded during time-lapse imaging. Scale bar, 100µm. **c** Representative sequence of images during time-lapse acquisitions showing GFP+ migrating INs. Arrowheads show the displacement of control (white) and *Dyrk1a* haploinsufficient (orange) cell bodies. Scale bar, 50µm. **d-e** Quantification of the mean velocity (**d**) and the pausing time (**e**) of control and *Dyrk1a* haploinsufficient migrating INs. n= 15 (control), n= 24 (*Dyrk1a^DlxCre/+^*) explants from embryos of at least three different mothers, unpaired t-test, **p < 0.01, ****p < 0.0001. **f** Schematic representation of nucleokinesis and of the morphological and dynamic changes INs go through during migration. The leading process extends and branches toward the direction of migration. Then, the nucleus will translocate forward in a movement known as nucleokinesis (inside of a dashed box). Finally, the trailing process will retract. **g-h** The frequency (**g**) and amplitude (**h**) of the nucleokinesis were quantified in control and *Dyrk1a* haploinsufficient INs. n= 15 (control), n= 24 (*Dyrk1a^DlxCre/+^*) explants from embryos of at least three different mothers, unpaired t-test, ***p < 0.001. **i** Neurons were classified in four different categories according to their morphology between two nucleokinesis. Histogram shows the percentage of INs with each morphology. n= 12 (control), n= 12 (*Dyrk1a^DlxCre/+^*) explants from embryos of at least three different mothers, two-way ANOVA, **p < 0.01, ***p < 0.001. Data is represented like mean ± s.e.m. CGE, caudal ganglionic eminence

Efficient tangential migration also relies on the different morphological changes that INs go through during locomotion. This includes the extension and branching of a leading process and the retraction of the trailing process ^51,53^ (Fig. 3f). To further understand why IN positioning is disrupted, we studied morphological parameters of migrating INs. Besides the nucleokinesis defects, we noticed that the morphology of *Dyrk1a* haploinsufficient migrating INs was affected in both MGE and CGE-derived INs (Fig. 3i, Supplementary Fig. 4f-i, k-m) Compared to the control, these neurons showed abnormal branching of the leading process, with a significant increase in the number of INs with multiple leading processes. This increase was in detriment of the most commonly found morphology, the bifurcated single leading process (Fig. 3i, Supplementary Fig. 4f). Though branching of migrating INs was altered, the overall length of the segments remained unchanged (Supplementary Fig. 4g, k), and we did not detect any defect neither in the extension of the leading process nor in the retraction of the trailing process (Supplementary Fig. 4h-i, l-m).

Altogether, these observations indicate that *Dyrk1a* promotes tangential migration of cortical INs cell-autonomously, by regulating velocity, nucleokinesis and controlling neurite branching during tangential migration.

### Tangential migration relies on DYRK1A kinase activity

DYRK1A is a serine-threonine-tyrosine kinase, implicated in the regulation of many cellular mechanisms ^20,21^. Our results clearly showed the importance of the genetic dosage of this gene for the correct migration of INs within the cortex. However, how and to what extent DYRK1A’s kinase activity is involved in the regulation of this process is still not known. To tackle this question, we generated a new knock-in mouse model for *Dyrk1a* carrying the K188R kinase dead mutation (*Dyrk1a^K^*^188*R/+*^ mice) (Fig. 4a) ^36,54^. Heterozygous mice show reduced DYRK1A kinase activity, with activity levels reaching around 65% compared to those of the controls (Fig. 4b). *Dyrk1a^K^*^188*R/+*^ mice were crossed with the GAD65-EGFP mouse line, and both the distribution of INs within the developing cortex and the dynamics of migration of these cells were analyzed. We observed that, as in the case of haploinsufficiency, in INs with reduced kinase activity but where the copy number of *Dyrk1a* gene is not altered, GFP+ INs failed to efficiently enter the developing cortex (Fig. 4c, d). However, *Dyrk1a*^K188R/+^ brains also show an increase in both proliferation of IN progenitors and cell death in proliferative regions (Supplementary Fig. 5). To ascertain the cell-autonomous effect of reduced DYRK1A kinase activity in INs locomotion, we went back to the GEs explant culture set-up, where heterozygous kinase-dead INs will migrate on top of wild-type cortical feeders. We found that INs from *Dyrk1a*^K188R/+^ brains show similar defects in the migration dynamics to those we previously observed in INs lacking one copy of *Dyrk1a*, with a decrease in locomotion, associated to strong defects in nucleokinesis (Fig. 4e). To further validate the effect of DYRK1A kinase activity on INs migration, we treated wild-type GEs explant cultures with Leucettinib-21 (LCTB-21), a selective inhibitor of DYRK1A kinase activity ^55,56^. We found that 1h treatment with this inhibitor slows down the migration of INs compared to those treated with the kinase inactive isomer iso-Leucettinib-21 (iso-LCTB-21) as control (Fig. 4f). Together, this not only highlights the importance of *Dyrk1a* dosage but also of its kinase activity in the migration of cortical INs.

**Figure 4:**
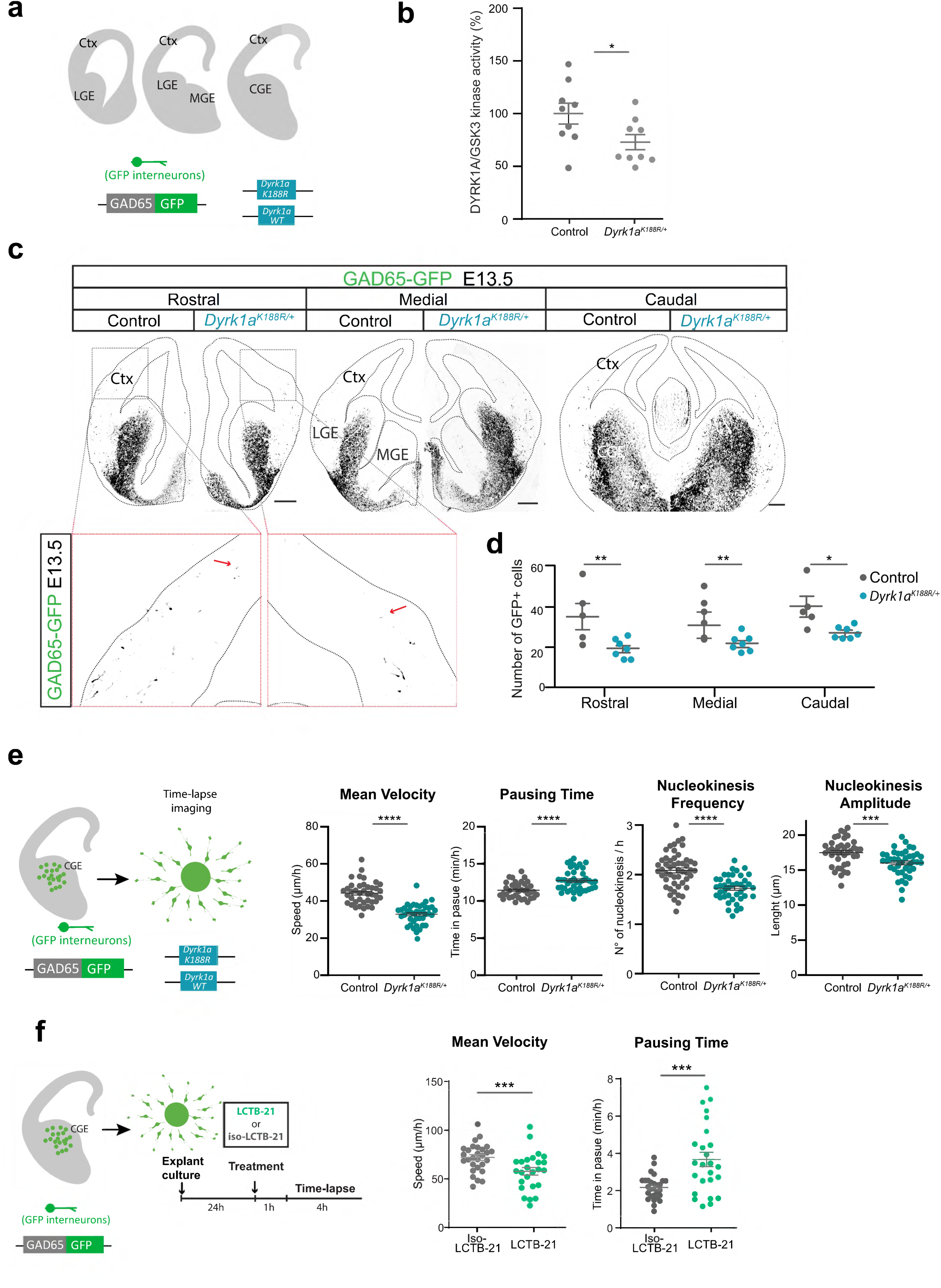
Tangential migration relies on DYRK1A Kinase activity. **a** Schematic representation of the three different E13.5 rostro-caudal sections of the brain analyzed. The importance of DYRK1A kinase activity the development of cortical INs was studied using a knock-in mouse model for *Dyrk1a* carrying the K188R mutation. INs were visualized thanks to the expression of the GFP under the GAD65 promoter. **b** *Dyrk1a^K^*^188^*^R/+^* mice have reduced DYRK1A catalytic activity. DYRK1A was purified from brain extracts by immunoprecipitation and GSK-3α/β was purified by affinity chromatography on axin-agarose beads. Activities of the purified kinases were assayed in triplicate in a radioactive kinase assay using specific peptide substrates, and are reported after normalization with wild-type GSK-3α/β activities. n= 9 brains (control and *Dyrk1a^K^*^188^*^R/+^*), unpaired t-test, *p < 0.1. **c** Representative brain sections of control and *Dyrk1a^k^*^188^*^R/+^* embryos. GFP-expressing INs (black) migrate from the GEs into the Ctx. The contour of the brain is delimited by a black dashed line. Higher magnification of the region inside the red dashed line. Red arrow shows the migration front. Scale bar, 200µm. **d** Quantification of the number of GFP+ INs entering the developing cortex in control and *Dyrk1a^K^*^188^*^R/+^* brains, at three different rostro-caudal levels. n=5 embryos (control), n=7 embryos (*Dyrk1a^K^*^188^*^R/+^*) from at least three different mothers, two-way ANOVA, *p < 0.05, **p < 0.01, ***p < 0.001. **e** Caudal ganglionic eminences explants of E13.5 control and *Dyrk1a^K188R/+^* embryos were cultured on top of wild type cortical feeders. One day later GFP+ INs exiting the explants were imaged for 5 hours (left panel). Quantification of the mean velocity, pausing time, and the frequency and amplitude of nucleokinesis of control and *Dyrk1a^K188R/+^*migrating INs (right panel) n= 40 (control), n= 41 (*Dyrk1a^K188R/+^*) explants from n= 4 (control), n=3 (*Dyrk1a^K188R/+^*) embryos from at least three different mothers, unpaired t-test, ***p < 0.001, ****p < 0.0001. **f** Caudal ganglionic eminences explants of E13.5 wild-type embryos were cultured on top of a synthetic coating and treated with 0.5µM leucettinib-21 or isoleucettinib-21. After one hour of treatment, GFP+ INs exiting the explants were imaged for 4 hours (left panel). Quantification of the mean velocity and the pausing time of control and treated migrating INs. n= 28 (isoLCTB-21), n= 25 (LCTB-21) explants from 4 embryos of three different mothers, unpaired t-test, ***p < 0.001 (right panel). Data is represented like dot plots with mean ± s.e.m. Ctx, cortex; LGE, lateral ganglionic eminence; MGE, medial ganglionic eminence; CGE, caudal ganglionic eminence; Iso-Lctb-21, isoleucettinib-21 ; Lctb-21, leucettinib-21

### *Dyrk1a* regulates actomyosin contractions during interneurons migration

Based on the cellular phenotype observed during INs migration, we analyzed gene expression in E15.5 *Dyrk1a^Dlxcre/+^* forebrains and compared it to control littermates. The analysis of *Dyrk1a^Dlxcre/+^* forebrains unraveled 1935 coding genes and 3052 predicted misregulated genes with an absolute log2 fold change superior to 0.5 and a corrected log P value superior to 2 (Supplementary Fig. 6a). Similarly, 1503 coding genes and 1650 predicted genes were differentially expressed in *Dyrk1a^+/-^* forebrains (Supplementary Fig. 6a). Investigating the protein-protein interaction landscape from *Dyrk1a^Dlxcre/+^* dysregulated genes highlighted enrichment in genes coding for G-coupled proteins. We were particularly interested in RHOA, associated with cytoskeleton regulation, mostly actin stress fibers formation and actomyosin contractility (Supplementary Fig. 6b). Then, looking at the expression of genes involved in cytoskeleton remodeling and nucleokinesis in both mutants ^57,58^, we found them more significantly downregulated in the *Dyrk1a^DlxCre/+^*mutant forebrains (Table S1; Supplementary Fig. 6c,d). Thus, we focused our efforts on understanding the implication of cytoskeleton remodeling in *Dyrk1a* migration defects. Indeed, the regulation of the actin cytoskeleton is critical for neural migration, and both nucleokinesis and neurite branching (which are both affected in *Dyrk1a* haploinsufficient and kinase-dead INs) are highly dependent on the actin cytoskeleton ^51,59,60^. Actin regulatory pathways, including PAK1 and cofilin, have been shown to be altered in GABAergic INs derived from DS patients’ iPSCs ^61^. Therefore, we tested whether the downregulation of DYRK1A could somehow affect these pathways. For this purpose, we performed biochemical analysis from isolated E14.5 GE of *Dyrk1a^DlxCre/+^* embryos. We observed that basal levels of the actin severing protein cofilin were not modified. However, in *Dyrk1a^DlxCre/+^*extracts, the phosphorylation state of cofilin was decreased, suggesting its activity is altered (Fig. 5a) . We further analyzed whether decreased cofilin activity could alter actin stability in *Dyrk1a* haploinsufficient cells. For this we performed an actin sedimentation assay, which allows us to separate soluble G-actin from filamentous F-actin. First, we observed that DYRK1A is mostly associated with actin fibers rather than to the soluble fraction of actin (Fig. 5b, c). Then, we found that the partial inactivation of *Dyrk1a* does not significantly modify the G/F actin ratio, neither in *Dyrk1a^DlxCre/+^* GEs (Fig. 5b) nor in mouse embryonic fibroblasts (MEF) derived from *Dyrk1a^+/-^* heterozygote full knock-out mice (Fig. 5c), suggesting that faulty cofilin activity observed in *Dyrk1a* haploinsufficient INs does not affect actin stability. However, biochemical analysis of GE extracts showed that *Dyrk1a* downregulation leads to decreased levels of Myosin light chain II (MLC-II) phosphorylation without affecting MLC-II levels (Fig. 5d) . As this phosphorylation induces MLC interaction with actin fibers, which results in the actomyosin contractions at the rear of the nucleus driving nuclear translocation in migrating INs ^60^, this could suggest decreased contractile activity during the migration of *Dyrk1a* haploinsufficient INs. To test this hypothesis, we monitored the dynamic rearrangement of the actin cytoskeleton during migration by expressing LifeAct-Ruby, which allows the visualization of F-actin without interfering with its dynamics ^62^ (Fig. 5e) . We performed live imaging of migrating INs, and observed that in the soma of control INs, low levels of F-actin are homogeneously distributed during phases of nuclear pause. Prior to nucleokinesis F-actin transiently condensates at the rear of the nucleus, and then goes back to initial levels once nuclear translocation is completed. Although the intensity of actin accumulation at the back of the nucleus just before nucleokinesis was similar to control (Fig. 5f), the timing of recruitment of F-actin in haploinsufficient INs was altered. In *Dyrk1a^DlxCre/+^*INs, the time between the beginning of actin recruitment and the actual nucleokinesis is longer (Fig. 5g) . Furthermore, the actin contraction area behind the nuclei is larger (Fig. 5h), which is generally associated with less efficient nucleokinesis ^63^. In line with this observation, nucleokinesis in *Dyrk1a* haploinsufficient INs took longer to complete (Fig. 5i). Finally, once nucleokinesis concludes, the dispersion of actin accumulation is less efficient, as it takes significantly more time to reach the initial basal intensity (Fig. 5j) . Together, our results show that *Dyrk1a* likely regulates IN migration through the modulation of actin contractibility.

**Figure 5:**
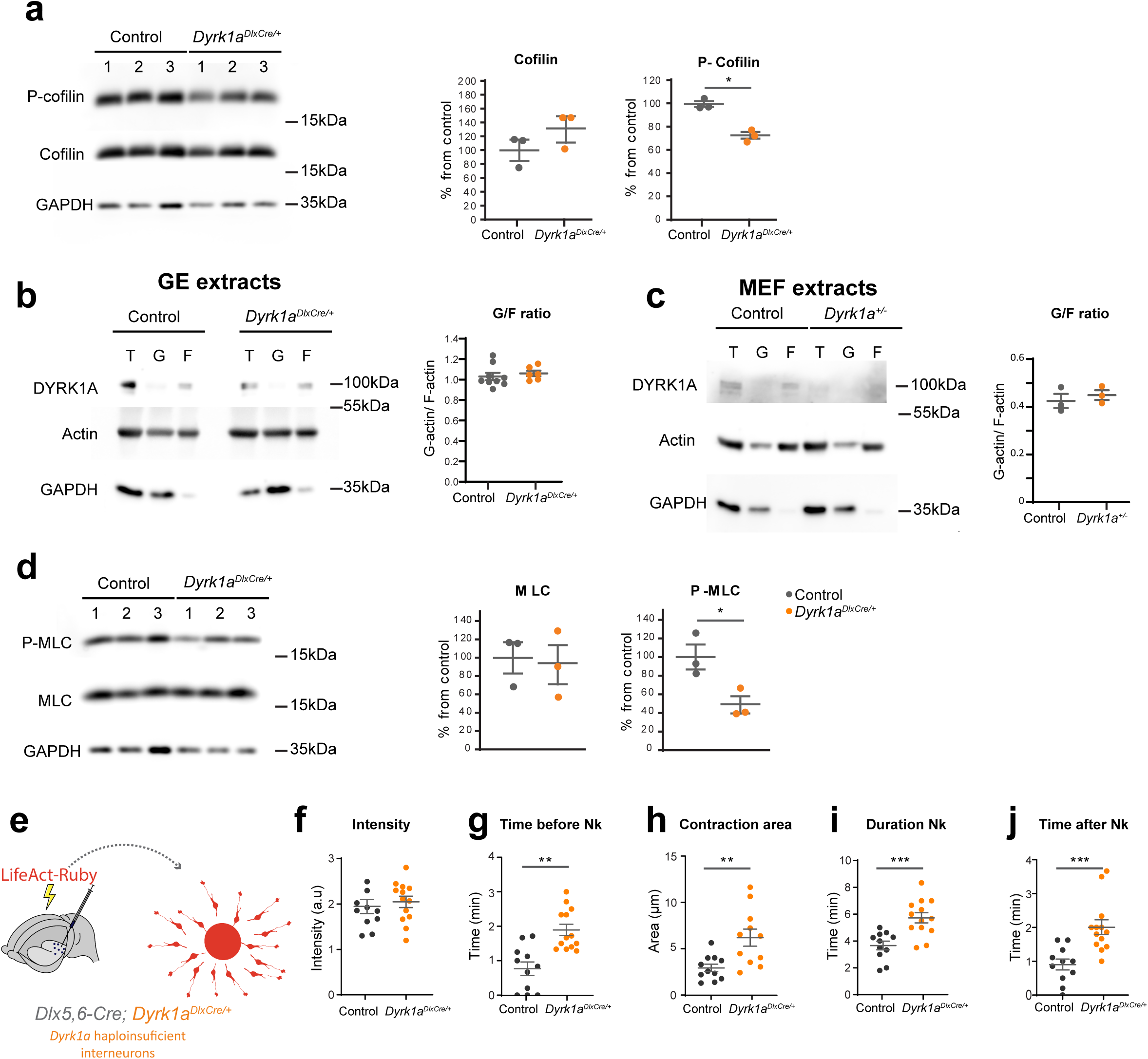
*Dyrk1a* regulates actomyosin contractions during INs migration. **a** Left: representative immunoblots showing cofilin expression and phosphorylation levels on protein extracts from E14.5 ganglionic eminences. Right: densitometric quantification of immunoblots against cofilin and phospho-cofilin. Relative intensities are expressed as percentage from the control. n= 3 (control and *Dyrk1a^DlxCre/+^*) embryos, unpaired t-test, *p < 0.1. **b** Left: Representative immunoblot of actin sedimentation assay, showing the expression of G- and F-actin and the distribution of DYRK1A within these fractions in extracts from E14.5 ganglionic eminences. Right: The ratio between G-actin and F-actin was quantified after densitometric quantification of actin bands. n= 9 (control), n= 6 (*Dyrk1a^DlxCre/+^*) ganglionic eminences, unpaired t-test. **c** Left: Representative immunoblot of actin sedimentation assay, showing the expression of G- and F-actin and the distribution of DYRK1A within these fractions, in extracts from mouse embryonic fibroblasts derived from *Dyrk1a^+/-^* heterozygote full knock-out mice. Right: The ratio between G-actin and F-actin was quantified after densitometric quantification of actin bands. n= 9 (control), n= 6 (*Dyrk1a^+/-^*) MEF cultures derived from different embryos, unpaired t-test. **d** Left: representative immunoblots showing MLC-II expression levels and phosphorylation on protein extracts from E14.5 ganglionic eminences. Right: densitometric quantification of immunoblots against MLC-II and phospho-MLC-II. Relative intensities are expressed as percentage from the control. n= 3 (control and *Dyrk1a^DlxCre/+^*) embryos, unpaired t-test, *p < 0.1. **e** E13.5 control and *Dyrk1a^DlxCre/+^* embryos were electroporated *ex-vivo* with LifeAct-Ruby, which stains F-actin. Time-lapse imaging was performed on electroporated ganglionic eminences explants to monitor actin dynamics during INs migration. (**f-j**) Quantification of the LifeAct-Ruby relative intensity at the rear of the nucleus (**f**), the time of actin recruitment before nucleokinesis (**g**), the size of the contraction area at the back of the nucleus during nucleokinesis (**h**), the duration of nucleokinesis (**i**) and the time of actin dispersion after these events (**j**) during INs migration. n= 10 (control), n= 13 (*Dyrk1a^DlxCre^*^/+^) cells from at least three different embryos and 3 different mothers. unpaired t-test, **p < 0.01, ***p < 0.001. Nk, nucleokinesis.

### *Dyrk1a* haploinsufficiency in interneurons leads to behavioral anomalies and epilepsy

Next, we wanted to understand whether and to what extent the specific downregulation of *Dyrk1a* in the GABAergic system affects behavior. As subjects with *DYRK1A* haploinsufficiency show motor delay and alterations in cognitive functions and social behavior, we analyzed if this correlates with abnormalities in *Dyrk1a^DlxCre/+^* adult mice.

We first focused on locomotor activity. We observed that, when placed in a novel environment, *Dyrk1a^DlxCre/+^* mice showed hyperactivity, with an increase in travelled distance in the Open Field (OF), in rearing in both the OF and elevated plus maze (EPM), and in the number of visited arms in the EPM and Y-maze (Fig. 6a, Supplementary Fig. 7b, d). In the OF, this increase in locomotion is observed during short periods (5min) and gradually declines with time (Fig. 6a). Hyperactivity was not found in the circadian test that lasted much longer and in which *Dyrk1a^DlxCre/+^* mice tend to have decreased locomotion and rearing (Supplementary Fig. 7a), suggesting that the hyperactivity behavior is linked to novelty exploration. Then, we analyzed the anxiety levels in these mice in the elevated-plus maze test, which revealed a decrease in anxiety levels as shown by the number of entries and time spent in open arms (Fig. 6b). Mice also showed less hesitation to enter open arms and explored the empty space more often (Supplementary Fig. 7b). However, this decreased anxiety was not observed in the percentage of time spent in the center of the OF (Fig. 6a).

**Figure 6:**
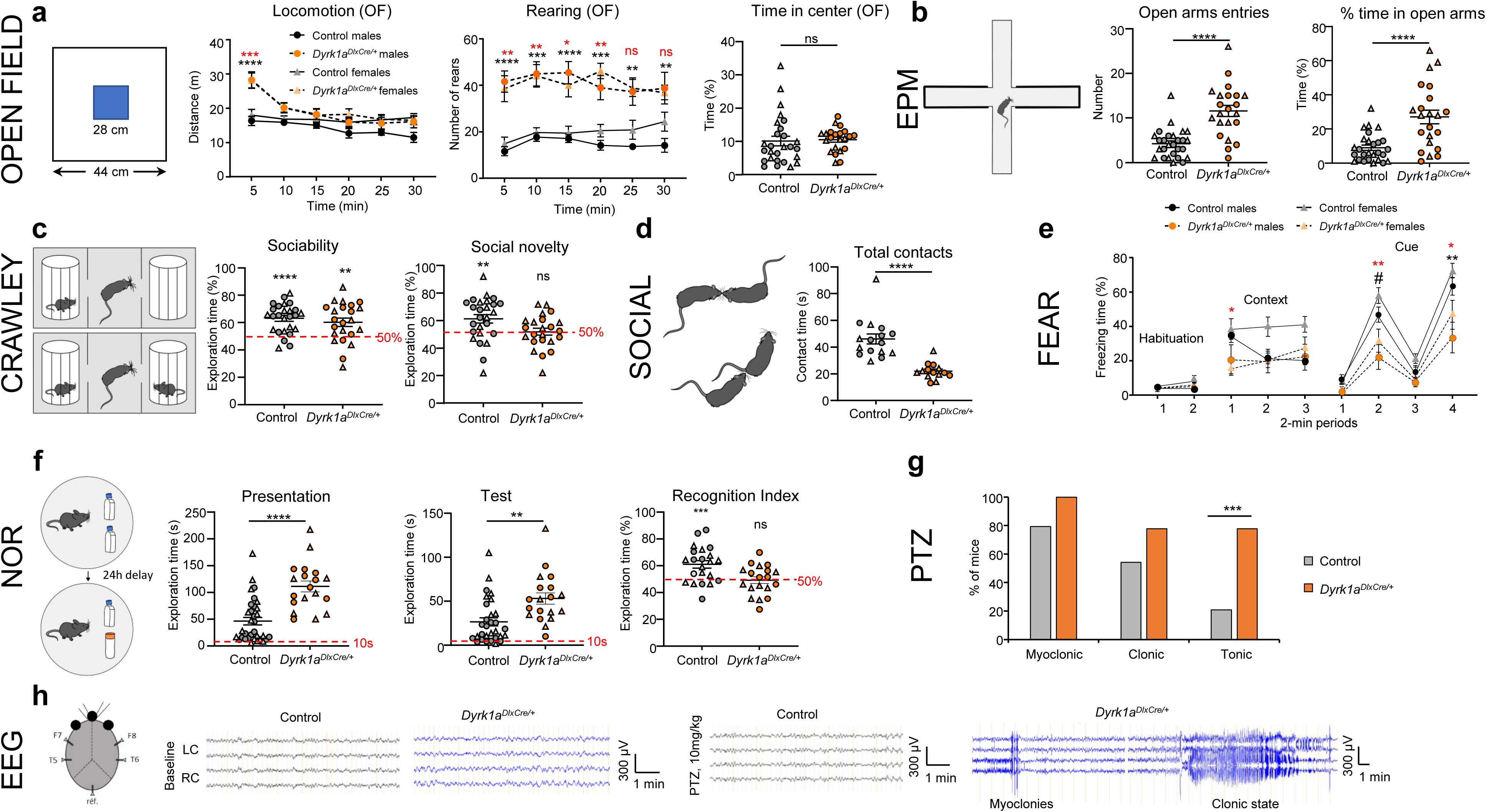
*Dyrk1a* haploinsufficiency leads to behavioral anomalies and epilepsy. **a** Locomotor assessment in the open field (OF) test. Schematic representation of the open field arena. The central area (blue) is used for assessing the emotional status of the mice. Distance travelled and number of rears per 5 minutes over the 30 min of the test, and mean percentage of time spent in the center area during the 30 min of the test (n = 14 males, and 13 females controls, n = 12 males and 10 females *Dyrk1a^DlxCre/+^*). Three-way repeated measure ANOVA with genotype and sex as between subject factor and sessions as within subject factor, with Tukey’s multiple comparisons test (for locomotion and rearing analysis). Two-way ANOVA with genotype and sex factors, *Dyrk1a^DlxCre^*^/+^ vs controls (time in center), *p < 0.05, **p < 0.01, ***p < 0.001, ****p < 0.0001. Black asterisk comparison between *Dyrk1a^DlxCre/+^*and control males and red asterisks for *Dyrk1a^DlxCre/+^* females vs control females. **b** Anxiety assessment in the elevated plus maze (EPM). Schematic representation of the EPM. Number of open arm entries and percentage time spent in the open arms (n = 14 males, and 13 females controls, n = 12 males and 10 females *Dyrk1a^DlxCre/+^*). Two-way ANOVA with genotype and sex factors, *Dyrk1a^DlxCre^*^/+^ vs controls, ****p < 0.0001. **c** Assessment of mouse sociability and preference for social novelty. Representation of the Crawley 3-chambers setup. Percentage of time exploring the cage containing the conspecific during the sociability test and percentage of time exploring the cage containing the new congener during the preference for social novelty test (n = 14 males, and 12 females controls, n = 12 males and 10 females *Dyrk1a^DlxCre/+^*). The percentage time of exploration is calculated from the total time exploring the two cages. One sample t-test vs 50% mean for each genotype, **p < 0.01, ****p < 0.0001. **d** Analysis of free social interaction. Representations of nose-to-nose and anogenital contacts that were used to assess social interaction. Total contact times recorded in pairs of mice (each dot plot represent a pair of mice from same genotype and of same sex (pairs of males are represented as circles and pairs of females as triangles). Two-way ANOVA with genotype and sex factors, *Dyrk1a^DlxCre^*^/+^ vs controls, ****p < 0.0001. **e** Fear conditioning analysis. Percentage of freezing per periods of two minutes during the habituation period (before conditional stimulus), the re-exposition to the same context 24 hours after conditioning and in the following cued session (precue: without conditional stimuli; cue: with conditional stimuli) (n = 21 males, and 21 females controls, n = 9 males and 11 females *Dyrk1a^DlxCre/+^*). Three-way repeated measure ANOVA with genotype and sex as between subject factor and sessions as within subject factor, with Tukey’s multiple comparisons post hoc analysis, #p<0.1; *p < 0.05, **p < 0.01. Black asterisks represent differences between control and *Dyrk1a^DlxCre/+^*males and red asterisks statistical differences between control and *Dyrk1a^DlxCre/+^* females. **f** Assessment of long-term declarative memory using the Novel Object Recognition (NOR) test. Schematic representation of the NOR paradigm. The retention time between presentation and test sessions was 24 hours. The type of object used in the presentation and the new object was randomly assigned for each mouse as well as its placement (left or right). Time spent exploring both objects during the presentation and tests sessions. Two-way ANOVA with genotype and sex factors, *Dyrk1a^DlxCre^*^/+^ vs controls, **p < 0.01, ****p < 0.0001. The recognition index corresponds to the percentage of time exploring the novel object in the test session (Novel/(Novel+Familiar)) with a value above 50% chance level indicative of memory performance (n = 9 males and 13 females controls, n = 10 males and 9 females *Dyrk1a^DlxCre/+^*). One sample t-test vs 50% mean for each genotype, ***p < 0.001. **g** Quantification of seizure susceptibility in *Dyrk1a^DlxCre^*^/+^ and controls mice after injection of 30 mg/kg of pentylenetetrazol (PTZ). Percentage of animals showing myoclonic, clonic and tonic seizures (n = 9 males and 15 females controls, n = 3 males and 6 females *Dyrk1a^Dlx-Cre/+^*). Chi square test, ***p < 0.001. **h** Electroencephalographic (EEG) analysis. Schematic representation of the mouse skull and the positions of the five electrodes. Representative EEG recordings from control and *Dyrk1a^DlxCre/+^*mice with, left, traces from left (Left Cortex, LC) and right (Right Cortex, RC) hemispheres of non-treated mice showing normal baseline cortical activity and right, traces from a PTZ-treated (10 mg/kg) control mouse showing normal activity and a *Dyrk1a^DlxCre/+^* mouse showing myoclonic and clonic activities. Data are represented as mean ± s.e.m. with single dots representing one animal. Females are represented with triangles and males with circles. Males and females are pooled in the same graph when the statistical analyses did not reveal significant effect of sex.

Then, as a proxy to study the autistic behavior observed in *DYRK1A* haploinsufficient patients, we assessed social interaction using the three-chamber test paradigm. In the first phase, when given the choice between an empty cage and a cage containing a never-encountered mouse, both control and *Dyrk1a^DlxCre/+^* mice spend more than 50% of the time exploring the cage containing the animal (Fig. 6c). However, in the second phase of the test, contrary to control animals, mutant mice show no preference for a new stranger mouse over the familiar one (presented in the first phase) (Fig. 6c), indicating reduced interest in social novelty. This finding was further supported by the deficit of social interaction observed in the free social test, shown by the net decrease in contacts of *Dyrk1a^DlxCre/+^* mice (Fig. 6d and Supplementary Fig. 7c).

We then subjected mice to classical learning and memory tests, to evaluate whether *Dyrk1a* inactivation in the GABAergic system affects cognition. First, to assess object memory, we performed the novel object recognition (NOR) task. In this test, though during presentation and retention phases mutant mice explored the objects significantly more than control mice, they showed no preference for the new object during retention phase, with a retention index not significantly different to 50% (Fig. 6f) . Thus, the mutant mice showed impairments in object memory and novelty discrimination. Next, we tested associative learning with the fear conditioning assay. *Dyrk1a^DlxCre/+^*mice displayed significantly lower contextual freezing during the first minutes. This was not due to the hyperactivity of the of *Dyrk1a^DlxCre/+^*mice as both control and mutant mice presented identical basal freezing levels before shock administration. The difference observed fades out with time, due to an increase in freezing time in *Dyrk1a^DlxCre/+^* females and a decrease of this parameter in control males. Cue specific freezing behavior was affected in both sexes, with a significant lower response to cue in *Dyrk1a^DlxCre/+^*mice (Fig. 6e). No defects in working memory were found in the Y-maze paradigm (Supplementary Fig. 7d), consistent with what was observed in the *Dyrk1a^+/-^* model ^27^ (Table S2). Together, this suggests that, contrary to working memory, long-term memory and associative learning depend on the correct dosage of *Dyrk1a* in the GABAergic system.

As epileptic seizures are a common comorbidity in *DYRK1A* haploinsufficient subjects ^12^, we analyzed the epileptic activity of control and *Dyrk1a^DlxCre/+^*mice. We first challenged mice with the pro-convulsive agent pentylenetetrazol (PTZ)(30mg/kg) and examined seizure susceptibility. We observed that in *Dyrk1a^DlxCre/+^* mice the occurrence of myoclonic, clonic or tonic seizures increased, reaching statistical significance at the tonic stage (χ2=12.7, p=0.0004) (Fig. 6g). Additionally, spontaneous generalized tonic-clonic seizures were detected in some *Dyrk1a^DlxCre/+^* animals (Movie 1) and never in control animals. Thus, to analyze these events, we performed a series of electroencephalogram (EEG) recordings. Baseline recording of untreated mice did not detect any abnormalities during the acquisition time. Therefore, we injected animals with 10 mg/kg of PTZ, which corresponds to a dose that is below the seizure threshold in control animals. Intraperitoneal PTZ injections resulted in abnormal EEG spikes and wave activity in 7 out of 10 *Dyrk1a^DlxCre/+^*mice followed by generalized tonic-clonic seizures that were observed in three *Dyrk1a^DlxCre/+^* mice (Fig. 6h). No epileptiform activity was recorded in the control mice (Fig. 6h). The increased susceptibility to epilepsy and cognitive deficits described correlate with what is observed in human subjects, highlighting the importance of the GABAergic dysfunction in *DYRK1A* haploinsufficiency pathogenesis.

## DISCUSSION

Studies on the essential role of DYRK1A in the correct development of the brain have been underway for a long time. Started with the *Drosophila* minibrain, *DYRK1A* was then investigated because of its location on human chromosome 21, involved in Down Syndrome (DS) – or Trisomy 21- and after 2012 with the *DYRK1A* haploinsufficiency syndrome (DHS) – due to the heterozygous loss of function of *DYRK1A*. However, most of these developmental studies have been biased toward understanding its role in the production of excitatory projecting neurons. Therefore, the specific role of *Dyrk1a* in the development of GABAergic inhibitory interneurons (INs) remained largely unclear.

In this study, we provide the first evidence of *Dyrk1a* as a key regulator in the migration of cortical INs. We demonstrate that migration dynamics is altered upon conditional loss of *Dyrk1a* or reduced DYRK1A kinase activity. More specifically, *Dyrk1a* controls the frequency of nucleokinesis and the amplitude of the forward progression of nuclei during these events, likely contributing to the net decrease of migration we observe in both mutants (*Dyrk1a^DlxCre/+^* and *Dyrk1a^K^*^188^*^R/+^*). Abnormalities in INs development have been commonly associated with neurodevelopmental disorders ^64^ and lately several studies have demonstrated that migration defects contribute to the etiology of conditions like ID and ASD ^65–69^. Reduced production of INs cell types and migration defects were also described in GABAergic neurons derived from DS iPSCs ^61^, and proposed as being partly responsible for the reduced number of INs observed in DS brains. Still, whether *Dyrk1a* overdosage is involved in this developmental defect is still to be uncovered. Here we show robust defects in INs migration *in vivo* 1) in *Dyrk1a* haploinsufficient INs derived from the MGE and CGE, 2) in INs with reduced DYRK1A kinase activity, and *in vitro* after pharmacological inhibition of DYRK1A kinase activity. Intriguingly, we show that in migrating pyramidal neurons, the heterozygous inactivation of *Dyrk1a* does not affect their positioning during corticogenesis. This rules-out cell-autonomous alterations in migrating glutamatergic neurons. One possible explanation is a lack of function and/or expression of *Dyrk1a* in this particular cell population. Even though the precise cellular and subcellular localization of *Dyrk1a* remains quite ambiguous, several studies suggest that *Dyrk1a* is not expressed in migrating pyramidal neurons. Pioneer studies of *Dyrk1a* spatio-temporal localization during brain development, show that *Dyrk1a* is downregulated in the intermediate zone of the developing cortex, where glial-guided migration of newborn pyramidal neurons occurs ^28^. More recent single-cell RNA sequencing (sc-RNAseq) shows that *Dyrk1a* mRNA is expressed in pyramidal progenitors and in mature neurons. However, expression levels in newborn migrating neurons seem to be negligible ^29^. These studies align with the concept that *Dyrk1a* expression decreases once progenitors exit the cell cycle to promote differentiation ^28,70^. In line with this idea, it has been shown that acute overexpression of wild-type *Dyrk1a* in pyramidal neurons leads to the accumulation of projecting neurons in the intermediate zone of the developing cortex, which could be a consequence of the ectopic expression of the protein in migrating neurons ^71^. It would therefore be relevant to study whether *Dyrk1a* overdosage, in the context of Down Syndrome, leads to glial-guided migration defects, or if expression control systems will avoid a pathological ectopic overexpression of *Dyrk1a* in these cells.

Interestingly, we proved that the kinase activity of DYRK1A is critical for the regulation of INs migration, with similar defects as those seen in *Dyrk1a* haploinsufficient INs. Reduced DYRK1A kinase activity additionally affects the proliferation of INs progenitors, with an increase of mitotic cells in proliferative regions of the ganglionic eminences. Primary microcephaly is one of the hallmarks of DHS ^10–18^ which is usually correlated with impaired neuronal production ^72,73^. Indeed, extensive studies performed in mice, zebrafish and fly have demonstrated that *Dyrk1a* controls cell cycle progression and cell cycle exit, through the regulation of DYRK1A expression levels. In neuronal precursors, *Dyrk1a* is transiently expressed promoting cell cycle exit and is then downregulated allowing prospective neurons to differentiate ^70^. In consequence, in pyramidal progenitors the downregulation of *Dyrk1a* leads to increased cell cycle, through mechanisms implicating Cyclin D1 and p27KIP1, modifying the output of excitatory neurons in the brain ^25,40,70,74^. It is therefore likely that DYRK1A kinase activity inhibition leads to an increased number of cells in proliferation as they are unable to exit cell cycle, and that this deregulation might ultimately lead to cell death ^70^. Whether this is the case for *Dyrk1a* haploinsufficient INs progenitors, and its consequences for cortical organization and function remains to be explored.

The migration of GABAergic INs is regulated by many extrinsic cues and intrinsic mechanisms, among which actin cytoskeleton reorganization has proven to be essential ^61,63,66,75,76^. During tangential migration actin condensates at the back of the nucleus, and actomyosin contractions generate the pushing forces that drive nucleokinesis ^60^. Here, we show that *Dyrk1a*-related nucleokinesis defects are associated with altered actomyosin dynamics during migration. First, we show an increase in the active form of the actin severing protein cofilin, which potentially results in increased actin turnover. However, despite the fact that cofilin is more active, we did not find any alteration in the soluble to fibrillary actin ratio (G/F ratio). This might be due to the fact that a decrease in cofilin phosphorylation may also result in increased recycling of actin filaments, or just the severing of actin filaments into smaller fragments ^77^. In either case, this will not be reflected in an absolute increase in the G/F ratio. Then we demonstrate that *Dyrk1a* haploinsufficiency leads to decreased phosphorylation of myosin light chain-2 (MLC-II). This phosphorylation is essential for MLC-II to generate the pushing forces needed for nucleokinesis, and uses actin filaments as the scaffold to generate these forces ^60^. Finally, by monitoring actin in *Dyrk1a* haploinsufficient migrating INs, we prove that actin dynamics at the rear of the nuclei prior to and during nucleokinesis is altered. Before nucleokinesis, F-actin accumulation rate is decreased, which probably leads to the net increase in pausing time observed during migration as nucleokinesis cannot occur. Then, during nucleokinesis, we observed an enlarged actomyosin contraction area in *Dyrk1a* haploinsufficient INs that leads to slower nucleokinesis ^63^. This is probably a conjunction of both less active MLC-II together to a decreased contraction efficiency due to decreased accessibility to long stable actin filaments. Altogether, these findings show that *Dyrk1a* controls tangential migration by regulating actin cytoskeleton organization during locomotion and contraction efficiency through MLC-II activation. Still, whether *Dyrk1a* directly interacts with cofilin and MLC-II or if its entry point is upstream in regulatory pathways is still to be investigated.

In addition to the developmental alterations, the specific haploinsufficiency of *Dyrk1a* in GABAergic inhibitory INs is sufficient to generate cognitive and social deficits in adult mice, mirroring clinical manifestations observed in DHS, such as ID and ASD ^10–18^. Previous findings have demonstrated that both germline *Dyrk1a^+/-^* mice, and heterozygous mice carrying a frame-shift mutation in *Dyrk1a* recapitulate these phenotypes observed in human ^32,34,36^. Still, the extent to which these findings are dependent on the GABAergic system remains unclear. Here, we demonstrate that *Dyrk1a* heterozygous inactivation in the GABAergic system leads to impairments in long-term memory and associative learning. These observations are supported by our previous work showing that the pathology-associated cognitive processes are not related to glutamatergic neurons, as no deficit in the NOR and fear conditioning tests were observed when removing one copy of *Dyrk1a* in pyramidal neurons postnatally ^27^. Nevertheless, this does not seem to be so clear-cut, as a complete lack of *Dyrk1a* in glutamatergic neurons results in stronger cognitive deficits, both in NOR and fear conditioning ^27^. Thus, whether *Dyrk1a* expression levels – low expression in pyramidal neurons is sufficient to fully perform its function - or expression timing - excision of *Dyrk1a* during the development of pyramidal neurons – are relevant for glutamatergic neuron physiology requires further investigation. Our data also show that *Dyrk1a^DlxCre/+^*mice exhibit reduced social preference and interactions, suggesting an important role of the GABAergic system in the pathogenesis of ASD-like phenotypes in DHS. Interestingly, similar social deficits have been documented in models affecting exclusively the excitatory system. Mice with heterozygous inactivation of *DYRK1A* in the developing excitatory neurons ^35^ and the full postnatal inactivation of *DYRK1A* in the mature pyramidal neurons (Brault et al., 2021), both lead to similar social deficits.

The disruption of the E/I balance in cortical circuits has been proposed as one of the key mechanisms underlying ASD ^78–80^. This can implicate developmental dysregulations and/or a functional imbalance of either the GABAergic or glutamatergic systems. So, this raises the question of whether social deficits in DHS are a result of an acquired E/I imbalance, regardless of its origin, or if deregulations in both GABA and glutamatergic systems will have synergistic effects, for example by lowering the balance level. On the other hand, epileptic seizures seem to be specifically related to the haploinsufficiency in the GABAergic system, resulting in spontaneous seizures and an increased susceptibility to PTZ-induced epileptic seizures. This aligns with previous studies indicating that pyramidal neurons are not implicated in *Dyrk1a*-associated seizures ^27^. The specific involvement of INs development and migration in epilepsy has also been shown, for example, in the X-linked aristaless-related homeobox (ARX). ARX mutations in human lead to ID and epilepsy and mouse models with targeted deletion of ARX develop epileptic seizures when the gene is deleted in GABAergic neurons and not in pyramidal neurons ^69,81,82^, further supporting a key role of the GABAergic system in epileptogenesis.

Altogether, our findings point to a central role of *Dyrk1a* dosage in the development of GABAergic INs, through the regulation of actomyosin during tangential migration. We also prove that the *Dyrk1a^DlxCre/+^* model is a relevant model to study the functional consequences of *Dyrk1a* downregulation in inhibitory INs, as it recapitulates several phenotypes seen in DHS patients. Our work also strongly justifies widening the investigations on the contribution of *Dyrk1a* in the development of the GABAergic system in the context of DS. *DYRK1A* is a key gene for therapeutic approach in DS ^21,55^ and knowing how its increase in gene dosage, and at which precise time during development or adulthood, contributes to the DS pathophysiology, will be key for considering new perspectives in therapeutic approaches.

## MATERIALS AND METHODS

### Experimental Model and genotyping

All experimental procedures were done in an authorized establishment (Institute Clinique de la Souris, license C67-218-40) and in agreement with the European Community Laboratory Animal Care and Use Regulations (2010/63/UE86/609/CEE) and the French ministry of Agriculture (law 87 848), which approved experimental protocols used (APAFIS agreements #15169 and #15691). Mice were housed in groups (2-5 animals) under a 12:12-hour light/dark cycle with constant temperature (22 ± 1°C) and humidity (55 ± 10 %), and with access to food and water *ad libitum.* CD1 mice were purchased from Charles Rivers (France), and used for the production of E13.5 embryos for cortical feeders. All other mouse models were maintained on C57BL/6NJ background. Gad65-EGFP transgenic mouse line ^43^ (kindly provided Gábor Szabó) expresses the GFP under the control of the GAD65 promoter, and was used to visualize GABAergic INs. The transgenic Tg(dlx5a-cre)1 ^Mekk^ (IMSR_JAX:008199) lines, named here DlxCre ^42^ were used to delete the targeted conditional knock-out allele specifically in GABAergic postmitotic INs. The transgenic Tg(mI56i-cre,EGFP)1^Kc^ (IMSR JAX:023724) line, named here Dlx-Cre-IRES-GFP ^52^, was used to delete the targeted conditional allele in GABAergic postmitotic INs and expressing ate the same time the GFP reporter. *Dyrk1a* conditional knock-out mice (*Dyrk1a^cKO/+^; Dyrk1a ^tm^*^1^*^.2Ics^*; MGI:6783456) containing lox sequences sites flanking exon 7 of the endogenous gene, was generated as previously described ^27^. The *Dyrk1a^K^*^188^*^R/+^* mouse line, which lacks of DYRK1A kinase activity, was generated using targeted recombination . The targeting vector was constructed as follows. A 3.7 kb genomic fragment encompassing *Dyrk1a* exons 5 and 6 was amplified by PCR (from BAC RP23-165H5) in two steps to allow the introduction of the A>G point mutation and subcloned in an ICS proprietary vector. This mutation changes the AAA codon in an AGA codon leading in a Lysine to Arginine mutation at position 188 (K188R). This ICS vector has a floxed Neomycin resistance cassette associated with a Cre-autoexcision transgene. A 3’ homology arm of 3 kb was amplified by PCR and subcloned in step1 plasmid to generate the final targeting construct. A crRNA AATGTGGCCTGGCTATAAT (MIT specificity score of 85 (http://crispor.tefor.net/) targeting the sequence at the site of insertion of the NeoR selection cassette was cloned in pX330 from Addgene (CRISPR/Cas9 plasmid #42230. Both plasmids (targeting and CRISPR/Cas9 plasmids) were electroporated circular in C57BL/6N mouse embryonic stem cells (ICS proprietary line). After G418 selection, targeted clones were identified by long-range PCRs and further confirmed by Southern blot with an internal (Neo) probe and a 5’ external probe. One positive ES clone was validated by karyotype spreading and microinjected into BALB/C blastocysts. Resulting male chimeras were bred with wild type females. Germline transmission with the direct excision of the selection cassette was achieved in the first litter.

All genotyping was performed by standard short-range polymerase chain reaction (PCR) with primers listed in Table S3. Both males and females were used for this study.

### Brain collection and sectioning

Embryos were killed by decapitation. After dissection, embryonic and brains were fixed over-night (ON) in 4% paraformaldehyde (PFA) at 4°C.

P8 pups were deeply anesthetized by an intraperitoneal injection of a solution containing 13 mg/kg xylazine (Rompun, 2%; Elanco) and 130 mg/kg ketamine (Ketamine 1000; Virbac) in NaCl 0.9%. Then, animals were intracardially perfused with PBS followed by perfusion with 4% PFA in PBS. Brains were then removed from the skull and post-fixed ON in 4% PFA at 4°C. Tissue was cryopreserved in a 20% sucrose solution in PBS at 4°C and then embedded in Cryomatrix (Thermo), frozen and stored at -80°C until further processing. Brains were then sectioned coronally at 14µm using a Cryostat (Leica), and kept at -80°C. For *in utero* electroporation analysis, after fixation, E18.5 brains were rinsed with PBS and embedded in 4% low-melt agarose (BioRad) diluted in PBS and were cut into 60μm-thick coronal sections using a vibrating-blade microtome (Leica VT1000S, Leica Microsystems).

### Brain sections, immunostaining, imaging and analysis

Cryostat brain sections were permeabilized and blocked in 2% normal goat serum (Sigma-Aldrich) in PBS-0.3% Triton X-100 (Sigma-Aldrich) for one hour, and then incubated ON at 4°C with the following primary antibodies in blocking solution: chicken anti-GFP (Invitrogen, 1:1000), rabbit anti-somatostatin (BMA-biomedials, 1:500), anti-neuropeptide-Y (Antibodies online, 1:500), rabbit anti-calbindin (Swant, 1:2000), rabbit phosphohistone-3 (Millipore, 1:500), rabbit anti-active caspase (R&D, 1:500). The corresponding Alexa-coupled secondary antibodies (Thermo Fischer) were incubated for 1 hour at 1:500. Nuclei were stained with a Hoechst solution 1:2000 (Life Technologies) for 20 min, and then mounted with Fluoromount-G mounting medium (Interchim). For somatostatin staining, prior to permeabilization antigen retrieval was performed using 10 mM tri-sodium citrate, 1.9mM citric acid, pH6.0, for 5 min at 95°C. Brain vibratome sections were processed for immunolabeling as follows: vibratome sections were permeabilized and blocked with 5% Normal Donkey Serum (NDS, Dominic Dutsher), 0.1% Triton X-100 in PBS. Slides were incubated with: chicken anti-GFP (Invitrogen, 1:800) antibody diluted in blocking solution overnight at 4 °C and secondary antibodies diluted in PBS-0.1% Triton one hour at room temperature, whereas cell nuclei were identified using DAPI (1 mg/mL Sigma). Slices were mounded in Aquapolymount mounting medium (Polysciences Inc).

For IN distribution and localization analyses, images were acquired using a time-lapse Axio observer microscope (Zeiss), equipped with a Hamamatsu OrcaFlash 4.0 camera using a Plan Aapochromat 20X objective for INs analysis. For pyramidal neurons studies, brains sections were images with a confocal microscope (Leica TCS SP8X equipped with a hybrid camera and a X20/0.70 objective) controlled by Leica Las X software. A Z stack of 1.5 μm was acquired in 512 ×512 mode and analyzed using ImageJ software (Java 1.8.0_172). Cell counting was done in at least four different brain slices of at least 8 different embryos for each condition. Only brains with comparative electroporated regions and efficiencies after histological examination were conserved for quantification. Cortical wall areas (upper cortical plate (Up CP), lower cortical plate (Lo CP), intermediate zone (IZ), subventricular zone (SVZ)/ventricular zone (VZ)) were identified according to cell density (nuclei staining with DAPI). The total number of GFP-positive cells in the embryonic brain sections was quantified by counting positive cells within a box of fixed size and the percentage of positive cells in each cortical area was calculated.

### Ganglionic eminences explant culture

Ganglionic eminence explants were cultured on top of wild type cortical feeders as detailed in ^83^. For this, a layer of wild type cortical neurons from E13.5 embryos were seeded on a glass bottom dish (MatTek or CELLView) previously coated sequentially with poly-L-Lysin (0.1mg/ml, Sigma-Aldrich) and laminin (10µg/ml, Sigma-Aldrich). Feeders were left to attach for one hour and then DMEM/F12 media supplemented with 2mM L-glutamine, 33mM D-Glucose, 1% N2, 2% B27 with vitamin A and 1% penicillin/streptomycin was slowly added. Cultures were maintained in an incubator at 37°C with 5% CO2. The following day, ganglionic eminences of the different GFP-expressing mouse lines were dissected and cut into 8-10 fragments. Media was removed from cortical cultures leaving a minimum volume, and explants were then placed on the top of the feeder layer and left to attach. One hour later, neurobasal medium supplemented with 1% L-glutamine, 33mM D-Glucose, 2% B27 containing vitamin A, 5mM pyruvate, 1% N2 and 1% penicillin/streptomycin was gently added and co-cultures were kept in the incubator for at least 12 for further processing.

Alternatively, for pharmacological treatment, ganglionic eminences explants were cultured on glass bottom petri dishes previously coated with poly-L-Lysin (2mg/ml, Sigma), fibronectin (5µg/cm ^2^, Sigma) and N-Cadherin (5µg/ml, R&D) ^84^.

### *Ex vivo* electroporation

For actin dynamic visualization, we used the pCAGGGS-lifeAct-Ruby plasmid (1µg/µl). Plasmids were prepared using a NucleoBond Xtra Maxi kit (Macherey-Nagel) and eletroporated *ex-vivo* as described by ^83^. E13.5 embryos were decapitated and ganglionic eminences exposed. Expression vectors combined with Fast Green (2µg/ml, Sigma) were then injected into the ganglionic eminences using pulled glass capillaries (Harvard Apparatus). Electroporation was done by placing 3mm platinum disk electrodes (Sonidel) on both sides of the embryonic brains, and delivering 5 electric pulses at 50V for 50ms at 950ms intervals using a CUY21EDIT electroporator (Sonidel). After electroporation GEs were micro-dissected and cultured on top of cortical feeders as described above.

### *In utero* electroporation

*In utero* electroporation was performed as previously described ^85^. Briefly, pregnant mice at 14.5 days post-coitus were anesthetized with isoflurane (2 L per min of oxygen, 4% isoflurane in the induction phase and 3% isoflurane during surgery operation; Tem Sega). The uterine horns were exposed, and a lateral ventricle of each embryo was injected using pulled glass capillaries (Harvard apparatus, 1.0 OD*0.58 ID*100 mmL) with Fast Green (1 μg/μL; Sigma) combined with 3 μg/μL NeuroD:Cre-IRES-GFP construct and 1 μg/μL NeuroD:IRES-GFP vector using a micro injector (Eppendorf Femto Jet). Plasmids were further electroporated into the neuronal progenitors adjacent to the ventricle by discharging five electric pulses (35 V) for 50 ms at 1 s intervals using electrodes (diameter 3 mm; Sonidel CUY650P3) and ECM-830 BTX square wave electroporator (VWR international). After electroporation, embryos were placed back in the abdominal cavity and the abdomen was sutured using surgical needle and thread. Pregnant mice were killed by cervical dislocation 4 days after surgery (E18.5) and embryos were collected.

### Time-lapse acquisition and analysis

For time-lapse videomicroscopy of INs migration, ganglionic eminences explant cultures were placed in a humidified and thermoregulated chamber maintained at 37 °C and 5% CO2 on the stage of a spinning-disk microscope (Leica), equipped with an Orca Flash 4.0 Camera. For INs migration studies, 16 to 18 successive 1µm z optical planes were acquired every 5 min during 5 h using a 20x objective. For analyzing actin dynamics during migration, we used a 40x HC PL APO oil objective. Images were acquired every 45 sec for three hours, performing 12 successive z optical planes (1µm) at every timepoint. Sequences were analyzed using the ‘Manual Tracking’ and ‘Track Mate’ Plugins in Image J.

### Explant culture immunostaining

Ganglionic eminences cultures were fixed in PFA 4% for 20 min, and then permeabilized and blocked in 2% normal goat serum (Sigma-Aldrich) in PBS-0.3% Triton X-100 (Sigma-Aldrich) for one hour. Primary antibodies were incubated ON at 4°C with the following primary antibodies in blocking solution: rabbit-anti-DYRK1A (Abcam, 1:100), mouse anti-DYRK1A (Abnova, 1:100), rabbit anti-myosin light chain 2 (Cell signaling, 1:100), mouse anti-phospho myosin light chain 2 (Cell signaling, 1:1000), rabbit anti-phospho myosin light chain 2 (Cell signaling, 1:1000), rabbit anti-cofilin (Abcam, 1:100) and rabbit anti-phopsho-cofilin (Cell signaling, 1:100).The corresponding Alexa-coupled secondary antibodies (Thermo Fischer) were incubated for 1 hour at 1:500 dilution in PBS. Nuclei were stained with a 1:2000 Hoechst solution (Life Technologies) for 20 min. Images were acquired using a confocal microscope TCS SP8 X (Leica microsystems).

### Pharmacological treatment

Ganglionic eminences explants cultures were treated with 0.5µM of LCTB-21 or isoLCTB-21. One hour before time-lapse imaging, compounds were diluted to 1µM in supplemented culture media. Half of the culture media was replaced by this solution and treatment was maintained throughout the acquisition.

### Actin assay

E15.5 GEs were dissected, and lysed in a buffer containing 50 mM PIPES pH 6.9, 50 mM NaCl, 5 mM MgCl2, 5 mM EGTA, 5% glycerol, 0.1% NP40, 0.1% Triton, 0.1% Tween20, 0.1% ß-mercaptoethanol, using a 25G needle at room temperature, and then incubated during 10min at 37°C. Nuclei were removed by centrifugation at 2000g at room temperature. To separate soluble and fibrillary actin, the supernatant was centrifuged in a *TLA100.4 fixed angle* rotor at 100000 g for 1h at 37°C in an *Optima* TL ultracentrifuge (Beckman). The supernatant containing globular actin was collected, and the supernatant pellet enriched in fibrillary actin was resuspended in water containing 10mM cytochalasin D. Western blot analysis were performed by loading equal volumes of each fraction.

### Kinase activity assay

The kinase catalytic activity of DYRK1A was evaluated in control and mutant adult brains (n=9 per genotype) as described before ^86^. Protein extracts were prepared from frozen half brains (n=9 per genotype), homogenized in lysis buffer (1.2 mL/half brain) using Precellys® homogenizer tubes. After centrifugation at 5,000 rpm for 2 x 15 s, 1 mg of brain extract was incubated with 2 µg DYRK1A (H00001859 M01, Interchim, France) or GSK3-β (MBS8508391, Emelca Bioscience, France) antibodies at 4°C for 1 h under gentle rotation. 20 µL of Protein G Agarose beads (Thermo Fisher Scientific, France), prepared in bead buffer (50 mM Tris pH 7.4, 5 mM NaF, 250 mM NaCl, 5 mM EDTA, 5 mM EGTA, 0.1% Nonidet P-40 and protease inhibitor cocktail from Roche, France), were then added to the mix and gently rotated at 4°C for 30 min. After a 1 min spin at 10,000 x g and removal of the supernatant, the pelleted immune complexes were washed three times with bead buffer, and a last time with Buffer C (60 mM β-glycerophosphate, 30 mM p-nitrophenolphosphate, 25 mM Mops pH 7.2, 5 mM EGTA, 15 mM MgCl2, 2 mM dithiothreitol, 0.1 mM Na Orthovanadate, 1 mM phenylphosphate, protease inhibitor cocktail). DYRK1A or GSK-3 immobilized on beads were assayed in buffer C with Woodtide (KKISGRLSPIMTEQ) (1.5 µg/assay) or GSK3-tide (YRRAAVPPSPSLSRHSSPHQpSED-EEE, where pS stands for phosphorylated serine) as substrates. Kinase activities for each enzyme were assayed in buffer A or B, with their corresponding substrates, in the presence of 15 µM ATP in a final volume of 30 µL. After 30 min incubation at 30°C, the reaction was stopped by harvesting, using a FilterMate harvester (Packard), onto P81 phosphocellulose papers (GE Healthcare) which were washed in 1% phosphoric acid. 20 µL of scintillation fluid were added and the incorporated radioactivity measured in a Packard counter. Blank values were subtracted and activities calculated as pmoles of phosphate incorporated during the 30 min incubation. Controls were performed with appropriate dilutions of dimethylsulfoxide (DMSO). Kinase activities were expressed in % of kinase activity in control littermates.

### Expression analysis in E15.5 forebrains

Forebrains from control and mutant mice (n=3 per genotypes) were microdissected at the age of 15.5 days post coitum (E15.5) from the *Dyrk1a^DlxCre/+^* and *Dyrk1a^+/-^* lines. Then RNA was extracted using Trizol and Qiagen RNeasy Minikit following the manufacturer protocol, and sent for single-end bulk RNA-Seq sequencing. The preparation of the libraries was done by the GenomEast platform, using the TruSeq Stranded Total RNA Sample Preparation Guide PN 15031048. Total RNA-Seq libraries were generated from 150-300 ng of total RNA using TruSeq Stranded Total RNA LT Sample Prep Kit with Ribo-Zero Gold (Illumina, San Diego, CA), according to manufacturer’s instructions. Following purification, the depleted RNA was fragmented into small pieces using divalent cations at 94oC for 2 minutes. Cleaved RNA fragments were then copied into first strand cDNA using reverse transcriptase and random primers followed by second strand cDNA synthesis using DNA Polymerase I and RNase H. Strand specificity was achieved by replacing dTTP with dUTP during second strand synthesis. The double stranded cDNA fragments were blunted using T4 DNA polymerase, Klenow DNA polymerase and T4 PNK. A single ’A’ nucleotide was added to the 3’ ends of the blunt DNA fragments using a Klenow fragment (3’ to 5’exo minus) enzyme. The cDNA fragments were ligated to double stranded adapters using T4 DNA Ligase. The ligated products were enriched by PCR amplification (30 sec at 98oC; [10 sec at 98oC, 30 sec at 60oC, 30 sec at 72oC] x 12 cycles; 5 min at 72oC). Surplus PCR primers were further removed by purification using AMPure XP beads (Beckman-Coulter, Villepinte, France) and the final cDNA libraries were checked for quality and quantified using capillary electrophoresis. The molecule extracted from the biological material were polyA RNA. The sequencing was performed by the platform using Illumina Hiseq 4000.

The alignment was performed on the mouse GRCm38 release 93 using HISAT2 ^87^ with the options -p 8 – dta, -t –summary-file –no-unal, --no-hd. Samtools version 1.9 (http://www.htslib.org/) was used to generate the bam file and index and the counts were generated using HTSeq-Count 0.9.1 ^88^. Using RseQC ^89^ we checked that the percentage of mapped reads distributed in each genomic feature followed an optimal distribution (exon, 5’UTR exon, 3’ UTR exon, Intron, Intergenic…). Then differential expression analysis was done using FCROS ^90^. Representation of protein-protein interaction network was obtained from mysregulated genes using Cytoscape ^91^.

### Protein extraction and Western Blot

Dissected embryonic brain structures were lysed in RIPA buffer (Tris HCl 1M, pH 7.7; NaCl 5M; EDTA 0.5M; Triton X-100) supplemented with protease inhibitors (Roche) and phosphatase inhibitors (Sigma Aldrich) using a 25G needle. Protein concentration was then assessed using Bio-Rad protein assay reagent (Biorad). Samples were denatured at 95°C for 10 min in loading buffer then resolved in 6%, 8% and 10% acrylamide gels. Proteins were transferred onto nitrocellulose membranes, which were then blocked in 5% non-fat milk in TBS buffer, 0.1% Tween. Membranes were incubated with mouse anti-DYRK1A (Abnova, 1:1000), rabbit anti-myosin light chain 2 (Cell signaling, 1:1000), rabbit anti-phospho myosin light chain 2 (Cell signaling, 1:1000), rabbit anti-cofilin (Abcam, 1:1000), rabbit anti-phopsho-cofilin (Cell signaling, 1:1000) and mouse anti-actin (Sigma, 1:150.000) primary antibodies ON at 4°C, followed by an incubation with the corresponding HRP-coupled secondary antibody. Chemiluminescence was revealed using a SuperSignal West Pico or Femto Kits, and detected using an Imager 600 (GE Healthcare).

### Behavioral analysis

Two to three cohorts of control and *Dyrk1a^DlxCre/+^*mice were used for behavioral analysis with an age range starting at 2.5 months for the first test up to 9 months for the last one, as described in Table S4. Both males and females were used.

Tests were conducted between 8:00 AM and 4:00 PM. Animals were transferred to the experimental room 30 min before each experimental test. Behavioral experimenters were blinded as to the genetic status of the animals. We tested for locomotor activity, general and anxiety related behavior (circadian activity, open field and elevated plus maze tests), memory (Y maze, NOR and fear conditioning), social behavior (Crawley 3-chamber test and social interaction) and susceptibility to epileptic seizures (PTZ injections). As a majority of the animals from the first three cohorts did not sufficiently explore the objects in the NOR test, this test was optimized (see supplementary Materials and Methods for details) and a new batch of animals was used for this test. Number and age of the animals for each test are given in Table S4.

All the standard operating procedures for behavioral phenotyping have been already described ^27^ and are detailed in the supplementary Materials and Methods.

### Electroencephalographic (EEG) recordings

EEG was done as previously described ^92^. Electroencephalographic implantation was done under general anesthesia (Propofol, Fentanyl, Domitor mixture 100 μl/10 g). *Dyrk1a ^DlxCre/+^* (n = 10) and Control mice (n = 10) aged 5-9 months were implanted with homemade stainless-steel wire electrodes (Phymep, France). For each mouse, five single-contact electrodes were placed over the left and right frontoparietal cortices. The electrodes were secured into the skull and soldered to a microconnector that was fixed to the skull by acrylic cement. An electrode over the surface of the cerebellum served as ground for all derivations. All mice were allowed to recover for a period of 1 week before EEG recordings. Freely moving mice were then recorded in their housing transparent cage and were connected to a recording system. EEG signals were amplified with a band-pass filter setting of 0.1–70 Hz with a 64-channel system (Coherence, Natus Neurowork EEG, Natus Medical Inc., San Carlos, CA, USA) and sampled at 256 Hz. Recordings were performed during 1 h for evidence of spontaneous convulsive seizure. At the end of the recording period, animals were injected intraperitoneally with a convulsive dose of 10 mg/kg of pentylenetetrazole (PTZ; Sigma-Aldrich, Co), a GABAA receptor antagonist, to evaluate seizure threshold. EEG was recorded during 1 hour after PTZ. Video EEGs were reviewed off line for electrographic seizures.

### Analysis and statistics

All experiments consisted of at least three independent replicates. Statistical analyses were done using GraphPad Prism (GraphPad Software, Inc.). All data are presented as mean± standard error of the mean (s.e.m). One-way ANOVA was performed for multiple comparisons followed by the appropriate post hoc tests, whereas unpaired two-tailed Student’s t-test was used for dual comparisons. For behavioral studies data was analyzed using a 2-way ANOVA or 3-way repeated measure ANOVA with sex and genotype as factors unless stated otherwise in the text. When no difference was found between sex, the p-value comparing the genotype factor was used and males and females were pooled in the graph. Otherwise, Tukey’s multiple comparisons post-hoc test was used to compare difference between genotypes for each sex. P<0.05 was considered significant with *P<0.05, **P<0.01, ***P<0.001, ****P<0.0001. All the statistical tests were two-sided. No statistical methods were used to predetermine sample sizes but sample sizes were estimated on the basis of previous experiments in the laboratory and are similar to those generally employed in the field. No randomization procedure was applied in this study. Data distribution was tested using the Shapiro-Wilk test and the homogeneity of the variance with the F test (for unpaired t-test) or Brown-Forsythe (for ANOVA). The investigator was blinded for data collection and analysis, except for immunoblot analysis.

## Acknowledgements

Gábor Szabó for providing the GAD65-EGFP mice.

Laurent Meijer and Emmanuel Deau for providing LCTB-21 inhibitor and isoLCTB-21.

ICS animal facility, in particular Sophie Brignon and Milan Herrmann for their involvement in the project.

All the staff of the IGBMC Platform of photonic microscopy, for their assistance in imaging experiments, and the IGBMC Genomeast platform, a member of the “France Génomique” consortium (ANR-10-INBS-0009), for the transcriptome studies.

This work was funded by grants from the Fondation Jérôme-Lejeune (A.D and M-V.H, JLJ Postdoctoral fellowship and V.B for JLJ project N° 1647), the French state funds through the Agence Nationale de la Recherche under the project PRC DYRK-DOWN ANR-18-CE16-0020 (Y.H.; J.D.G), and support from program Investissements d’Avenir labeled IdEx Unistra (ANR-10-IDEX-0002), a SFRI-STRAT’US project (ANR 20-SFRI-0012), EUR IMCBio (ANR-17-EURE-0023) to JDG and YH, INBS PHENOMIN (ANR-10-INBS-07

PHENOMIN) to YH, the ‘France Génomique’ consortium (ANR-10-INBS-0009) for the GenomeEast platform, and INSERM/CNRS and University of Strasbourg. P.T. and A.D. are, respectively, research assistants and research engineer at the University of Strasbourg. VN is an engineer assistant at the CNRS. VA is a PhD student at the ED414 Unistra. MdM and TLN were respectiveley a post-doct and PhD at the CERBM GIE. VB and YH are respectively CNRS researcher and Principal investigator. J.D.G. is an INSERM Principal investigator. The funders had no role in the study design, data collection and analysis, decision to publish, or preparation of the manuscript.

All the members of Y.H and J.G research groups for their feedback.

## Author contribution

M-V.H conceived and designed the experiments, performed and analyzed experiments, coordinated the study and wrote the manuscript. A.D conceived and designed the experiments, performed and analyzed experiments. V.A. performed and analyzed time-lapse experiments with LCTB-21 treatment, and designed cartoons. G.R performed and analyzed EEG studies. T-L.N performed the kinase activity assay. P.T. performed *in utero* electroporations. M.deM performed bulk RNA-Seq analyses and V.N further analyzed this data. M-C.B. developed and validated the *Dyrk1a^K^*^188^*^R/+^* mouse model. J.D.G. conceived and supervised *in utero* electroporation experiments. V.B. conceived, performed and analyzed behavioral studies. Y.H conceived, coordinated and supervised the study, and provided the financial support.

## Supplementary Figure legends

**Supplementary Figure 1: *Dyrk1a* haploinsufficiency does not affect interneuron progenitor’s proliferation and interneuron cell death.**

**a** *Dyrk1a* localization in migrating INs. Ganglionic eminences explant cultures were stained against *Dyrk1a* with two different antibodies (rabbit anti-DYRK1A, green; mouse anti-DYRK1A, magenta) and counterstained with DAPI (blue). Scale bar, 5µm.

**b** Representative E12.5 and E15.5 brain sections showing ganglionic eminences of control and *Dyrk1a^DlxCre/+^* embryos immunolabeled against the mitotic marker PH3 (red). Scale bar, 200 µm

**c** Quantification of the number of PH3+ cells per mm in the three different ganglionic eminences. n=6 embryos (control), n=6 embryos (*Dyrk1a^DlxCre/+^*) in LGE; n=7 embryos (control), n=6 embryos (*Dyrk1a^DlxCre/+^*) in MGE, n=3 embryos (control), n=3 embryos (*Dyrk1a^DlxCre/+^*) in CGE, from at least three different mothers; unpaired t-test.

**d** *Dyrk1a^DlxCre/+^*and control cortices (left panel) and ganglionic eminences (right panel) were stained for the apoptotic marker cleaved caspase (red). GAD65-GFP+ cells are shown in green. Arrowheads indicate caspase+ cells. Scale bar, 100µm.

**e** Quantification of the number of caspase+ cells per mm ^2^ in the three different ganglionic eminences. n=7 embryos (control), n=11 embryos (*Dyrk1a^DlxCre/+^*), from at least three different mothers, two-way ANOVA.

Data is represented as dot plots with mean ± s.e.m.

Ctx, cortex; LGE, lateral ganglionic eminence; MGE, medial ganglionic eminence; CGE, caudal ganglionic eminence; PH3, phospho-histone 3

**Supplementary Figure 2: Postmitotic pyramidal neurons migration is not altered by *Dyrk1a* haploinsufficiency**

**a** *In utero* electroporation was performed on E14.5 control and *Dyrk1a^cKO/+^* embryos with NeuroD:Cre-IRES-GFP construct leading to the specific deletion of one copy of *Dyrk1a* in postmitotic pyramidal neurons. Embryos were collected and analyzed at E18.5.

**b** Representative images of E18.5 cortices from control, *Dyrk1a^cKO/+^* and *Dyrk1a ^cKO/cKO^* embryos electroporated with the NeuroD: Cre-IRES-GFP construct. GFP is expressed in electroporated neurons. Scale bar, 100µm

**c** Quantification of the percentage of GFP-positive neurons present in the upper-CP, lower-CP, IZ or VZ/SVZ in the control and *Dyrk1a^cKO/+^* embryos. n= 8 embryos for each condition, two-way ANOVA.

**d** Quantification of the percentage of GFP-positive neurons present in the upper-CP, lower-CP, IZ or VZ/SVZ in the control and *Dyrk1a^cKO/cKO^*embryos. n= 6 embryos for control and n = 7 for *Dyrk1a^cKO/cKO^*, two-way ANOVA.

Data is represented as dot plots with mean ± s.e.m.

CP, cortical plate; IZ, intermediate zone; VZ/SVZ, ventricular zone/subventricular zone.

**Supplementary Figure 3: *Dyrk1a* haploinsufficiency leads to abnormal localization of interneurons after birth.**

**a** Top: Schematic representation of the three P8 rostro-caudal brain sections and the cortical regions analyzed. Bottom: histogram represents the percentage of calbindin-positive INs per bin in control and *Dyrk1a^DlxCre/+^* dorso-lateral cortices, at three different rostro-caudal levels. n=4 brains (control), n=4 brains (*Dyrk1a^DlxCre/+^*) from at least three different mothers, two-way ANOVA.

**b** Top: Schematic representation of the three P8 rostro-caudal brain sections and the cortical regions analyzed. Bottom: histogram represents the percentage of somatostatin-positive INs per bin in control and *Dyrk1a^DlxCre/+^* dorso-lateral cortices, at three different rostro-caudal levels. n=6 brains (control), n=5 brains (*Dyrk1a^DlxCre/+^*) from at least three different mothers, two-way ANOVA.

Data is represented as dot plots with mean ± s.e.m. P8, postnatal day 8; CB, calbindin; SST, somatostatin.

**Supplementary Figure 4: *Dyrk1a* dosage controls INs migration dynamics and morphology**

**a** Medial ganglionic eminences explants of E13.5 control and *Dyrk1a^DlxCreGFP/+^* embryos were cultured on top of wild type cortical feeders. One day later GFP+ INs exiting the explants were imaged every 5 min for 5 hours.

**b** Representative sequence of images during 2 hours of time-lapse acquisitions showing GFP+ migrating INs. Arrowheads show the displacement of control (white) and *Dyrk1a* haploinsufficient (orange) nucleus. Scale bar, 50µm.

**c-e** Quantification of the mean migration velocity (**c**) and frequency (**d**) and amplitude (**e**) of nucleokinesis in control and *Dyrk1a* haploinsufficient migrating INs. n= 16 (control), n= 25 (*Dyrk1a^DlxCreGFP/+^*) explants from n= 4 (control), n=5 (*Dyrk1a^DlxCreGFP/+^*) embryos of at least three different mothers, unpaired t-test, *p < 0.05, ****p < 0.0001.

**f** Quantification of the mean number of primary, secondary and tertiary neurites in control and *Dyrk1a* haploinsufficient migrating INs. n= 16 (control) and n= 25 (*Dyrk1a^DlxCreGFP/+^*) explants from n= 4 (control) and n= 5 (*Dyrk1a^DlxCreGFP/+^*) embryos of three different mothers, two-way ANOVA, ***p < 0.001.

**g-i** Quantification of the mean length of primary, secondary and tertiary neurites (**g**), and of the leading (**h**) and trailing processes (**i**) in control and *Dyrk1a* haploinsufficient migrating INs. n= 16 (control) and n= 25 (*Dyrk1a^DlxCreGFP/+^*) explants from n= 4 (control) and n= 5 (*Dyrk1a^DlxCreGFP/+^*) embryos of three different mothers, two-way ANOVA for neurites length and unpaired t-test for the leading and trailing processes length.

**j** Schematic representation of the experimental set-up and mouse model used for INs morphology analysis. Caudal ganglionic eminences were cultured on top of wild-type cortical feeders and GFP+ INs migrating out of the explants were analyzed.

**k-m** Quantification of the mean length of different grade neurites (**k**), and of the leading (**l**) and trailing processes (**m**) in control and *Dyrk1a* haploinsufficient migrating INs. n= 15 (control and *Dyrk1a^DlxCre/+^*) explants for neurites length; n= 15 (control), n= 21 (*Dyrk1a^DlxCre/+^*) explants for leading and trailing process, from embryos of at least three different mothers, unpaired t-test for leading and trailing process length; two-way ANOVA for neurites length.

Data is represented as dot plots with mean ± s.e.m, or like histograms with mean ± s.e.m.

CGE, caudal ganglionic eminences; LGE, lateral ganglionic eminences; MGE, medial ganglionic eminences; I, primary neurites; II, secondary neurites; III, tertiary neurites.

**Supplementary** Figure 5**: DYRK1A kinase activity regulates interneurons progenitors’ proliferation and cell death.**

**a** Schematic representation of the three different E13.5 rostro-caudal sections of the brain analyzed. The importance of DYRK1A kinase activity the proliferation and cell death of cortical INs was studied using a knock-in mouse model for *Dyrk1a* carrying the K188R mutation

**b** Proliferation study. Representative brain sections immunolabeled against the mitotic marker PH3 (cyan). Scale bar, 200µm.

**c** The number of PH3+ cells per mm in the VZ (upper panel) or mm ^2^ in the SVZ (lower panel) was quantified in the three different ganglionic eminences. n=5 embryos (control), n=6 embryos (*Dyrk1a^K^*^188^*^R/+^*) from at least three different mothers, two-way ANOVA. *p < 0.05, **p < 0.01, ***p < 0.001.

**d** *Dyrk1a^K^*^188^*^R/+^* and control sections were stained for the apoptotic marker cleaved caspase. The number of caspase+ cells per mm ^2^ was quantified. n=3 embryos (control and *Dyrk1a^K^*^188^*^R/+^*) from three different mothers, two-way ANOVA. *p < 0.05

Data is represented dot plots with mean ± s.e.m.

Ctx, cortex; LGE, lateral ganglionic eminence; MGE, medial ganglionic eminence; CGE, caudal ganglionic eminence; PH3, phospho-histone 3

**Supplementary Figure 6: Expression studies of the *Dyrk1a* full heterozygous mutant or the conditional *DlxCre (Dlx)* allele in the developing brain at E15.5 unraveled dysregulation of RhoA-dependent pathways.**

**a** Volcano plot representing the expressed genes in the E15.5 brain isolated from *Dyrk1a^Dlx/+^* and *Dyrk1a^+/-^* carriers and compared to controls.

**b** Protein-protein interaction network derived from dysregulated genes identified in (a) with an absolute fold change > 0.5 and a corrected -log(p-value) > 2.62 and a false discovery rate >0.01. Each protein is represented by a dot and those which belongs to GPCR are circled in red.

**c** Table showing enrichment in the GPCR ligand binding proteins.

**d** Focused analysis showing the dysregulation affecting DYRK1A, RHOA, ROCK1, other kinases with myosin light and heavy chain coding genes involved in neuronal development in the *Dyrk1a^Dlx/+^* and *Dyrk1a^+/-^* brain at E15.5 ^58^.

**Supplementary Figure 7: *Dyrk1a* haploinsufficiency alters general behavior and cognition**

**a** Locomotor and rearing activities measured during circadian home cage monitoring over light and dark cycle periods. (n = 14 males, and 13 females controls, n = 12 males and 8 females *Dyrk1a^DlxCre/+^*). Three-way repeated measure ANOVA with genotype and sex as between subject factor and sessions as within subject factor, with Tukey’s multiple comparisons post hoc analysis, *p < 0.05, **p < 0.01. Black asterisks represent differences between control and *Dyrk1a^DlxCre/+^*males and red asterisks between control and *Dyrk1a^DlxCre/+^* females.

**b** Number of visited arms (open and closed arms), rears, explorations of empty space and hesitations in the open arms of the elevated plus maze (EPM) (n = 14 males, and 13 females controls, n = 12 males and 10 females *Dyrk1a^DlxCre/+^*). One male and one female control were removed from the exploration and hesitations in open arms as they did not enter an empty arm during the session. Two-way ANOVA with genotype and sex factors. *Dyrk1a^DlxCre/+^* vs controls, *p < 0.05, **p < 0.01, ***p < 0.001, ****p < 0.0001. Black asterisks represent differences between control and *Dyrk1a^DlxCre/+^* males and red asterisks between control and *Dyrk1a^DlxCre/+^* females.

**c** Nose-to-nose and anogenital contacts during free social interactions. Contact time was recorded for pairs of mice (each dot represents a pair of mice from same genotype and of same sex (pairs of males are represented as round dots and pairs of females as triangles). Two-way ANOVA with genotype and sex factors, *Dyrk1a^DlxCre/+^* vs controls, ****p < 0.0001.

**d** Assessment of short-term working memory using the Y maze test. Schematic representation of the Y maze. Number of visited arms scored as an index of locomotor activity during the 5 minutes session (n = 18 males and 15 females controls, n = 12 males and 12 females *Dyrk1a^DlxCre/+^*). Percentage of spontaneous alternation (%SPA) was assessed from animals that made 10 or more arm entries (four males and one female control were excluded in %SPA analysis as they did not reach the 10 arms exploration criterium for this test). Two-way ANOVA with genotype and sex factors, *Dyrk1a^DlxCre/+^* vs controls, ****p < 0.0001.

Data are represented as mean ± s.e.m. with single dots representing one animal. Females are represented with triangles and males with circles. Males and females are pooled in the same graph when the statistical analyses did not reveal significant effect of sex.

