## Supplementary figures and images for "Interneuron migration defects during corticogenesis contribute to *Dyrk1a* haploinsufficiency syndrome pathogenesis via actomyosin dynamics deregulations"

### Supplementary fig.1

#### Supplementary Fig. 1

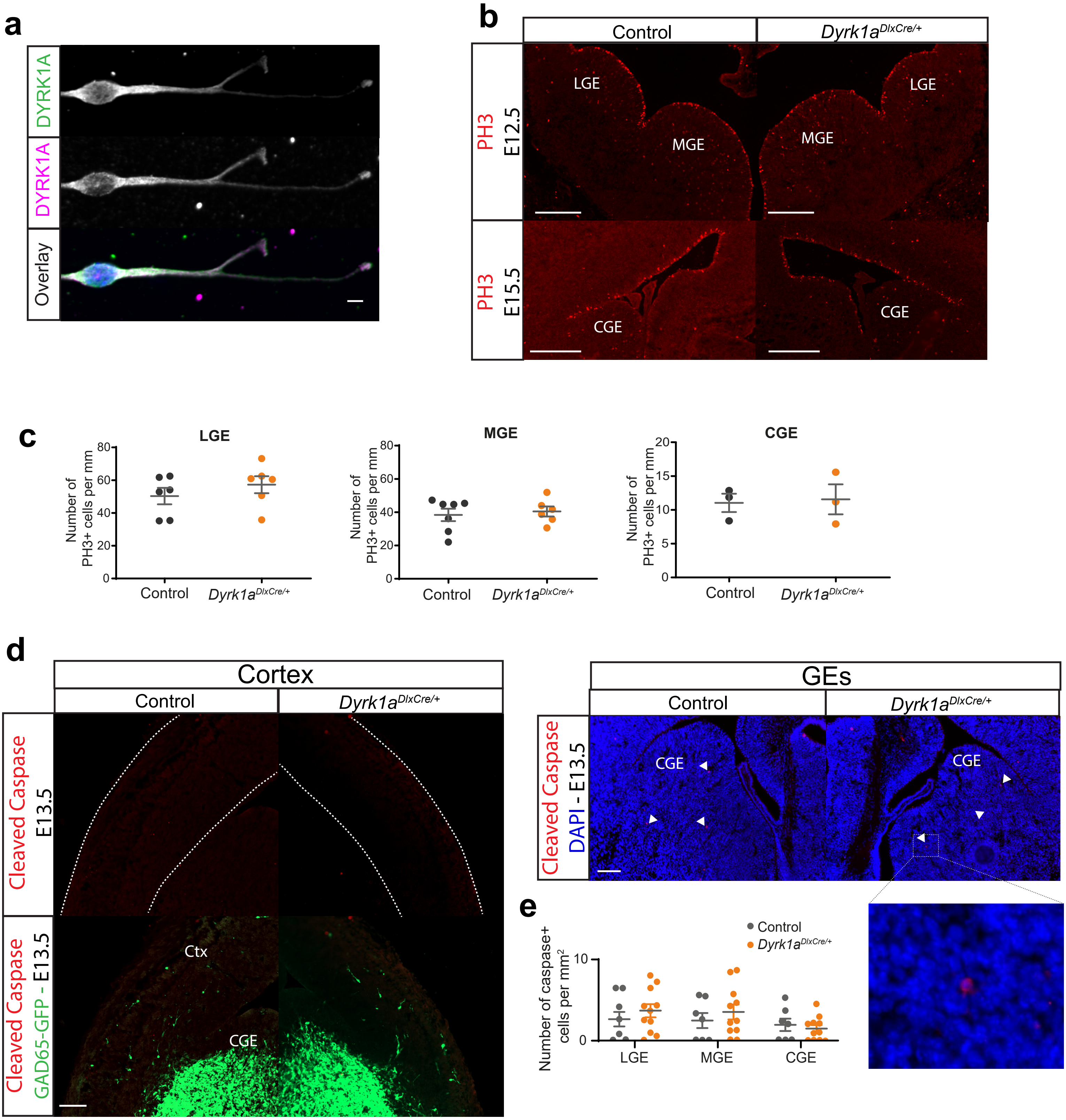

### Supplementary fig.2

#### Supplementary Figure 2

**a**

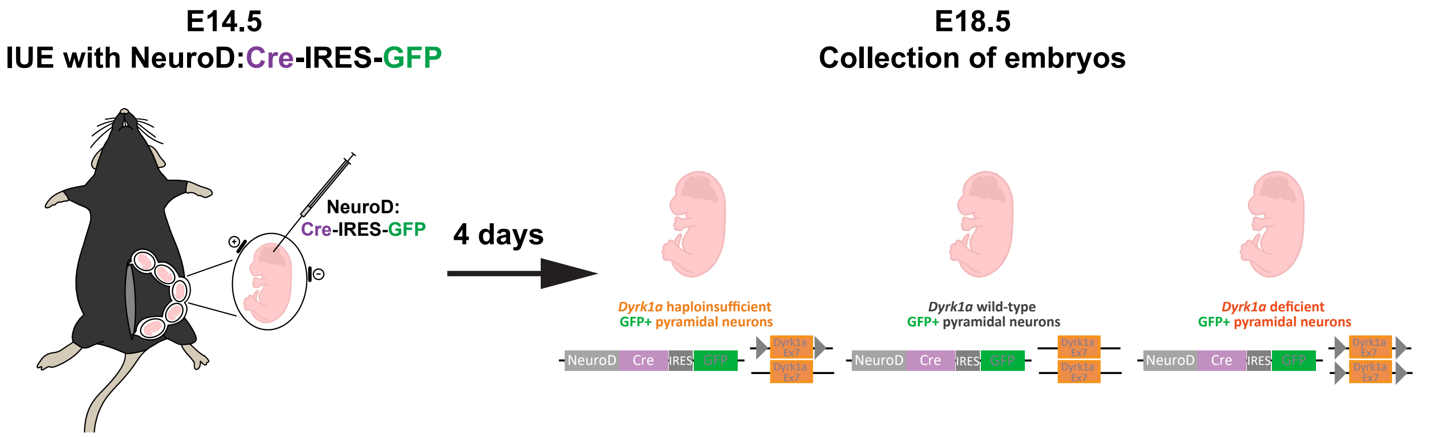


**b**

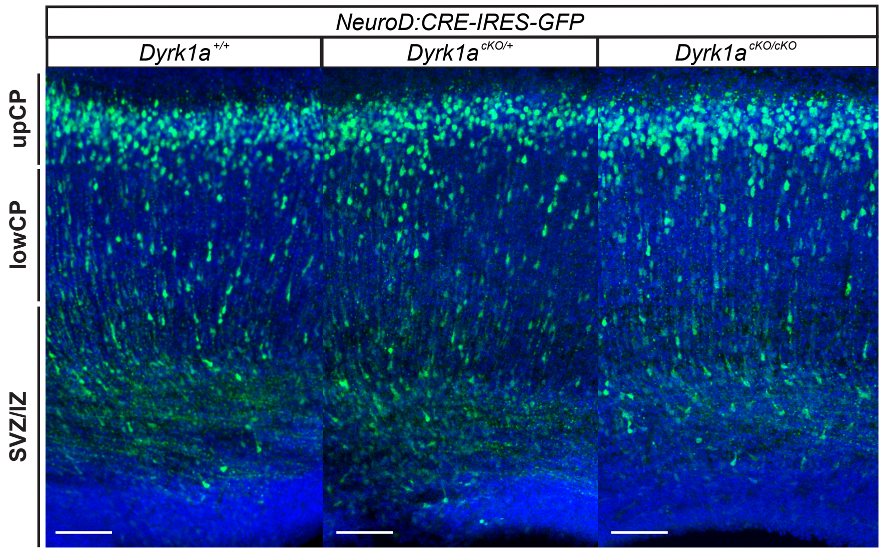


**c**

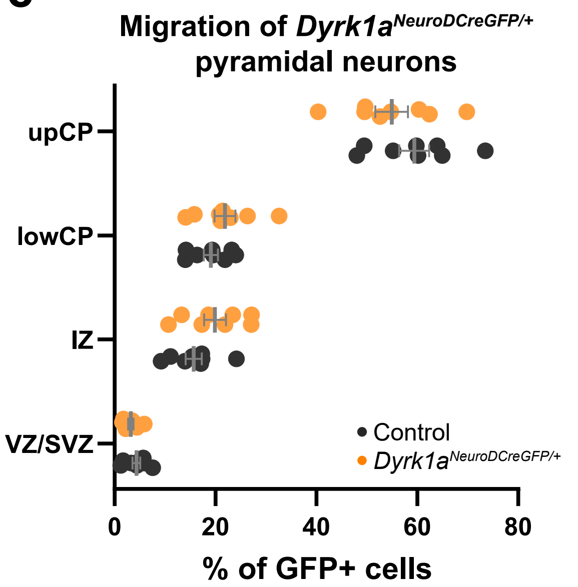


**d**

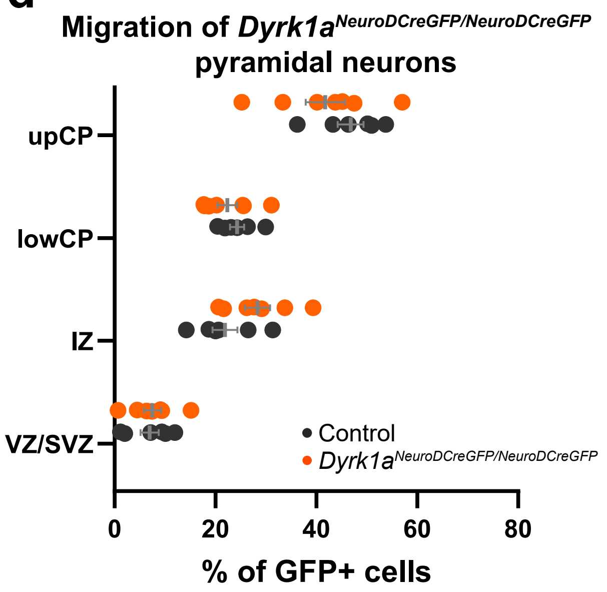

### Supplementary fig.3

#### Suplementary Fig. 3

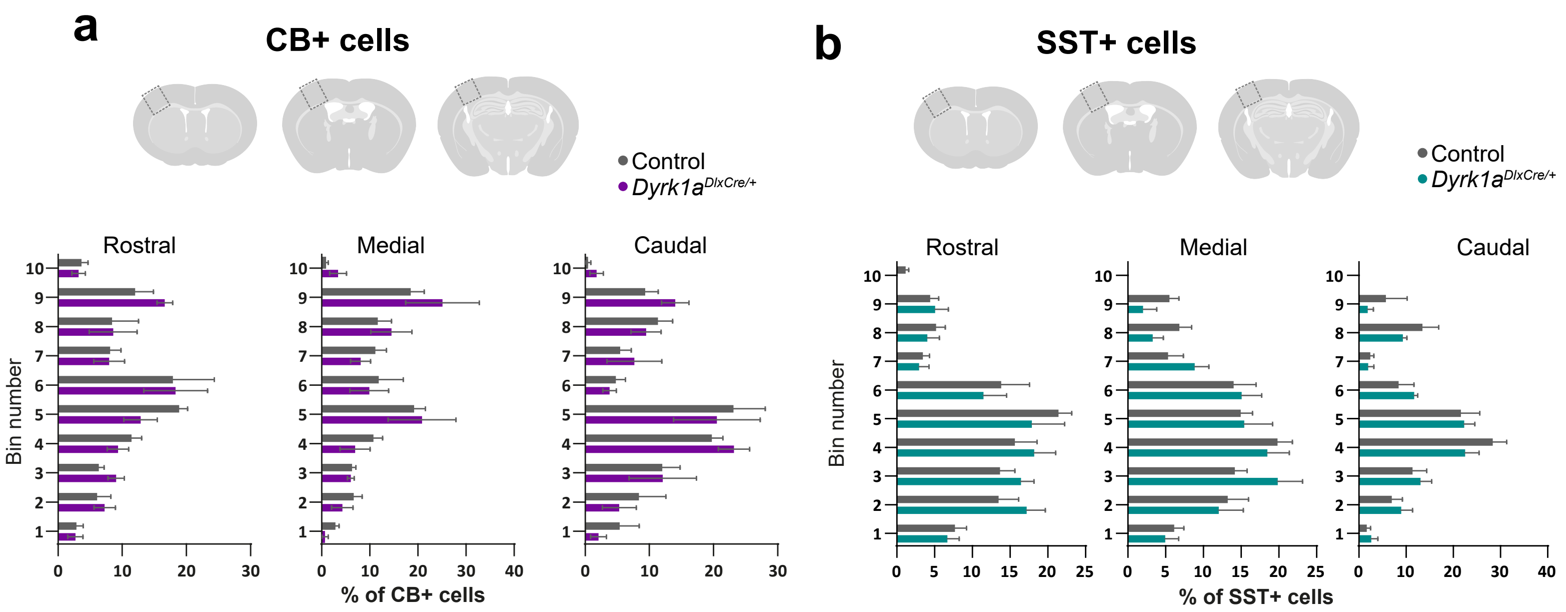

### Supplementary fig.4

#### Supplementary Figure 4

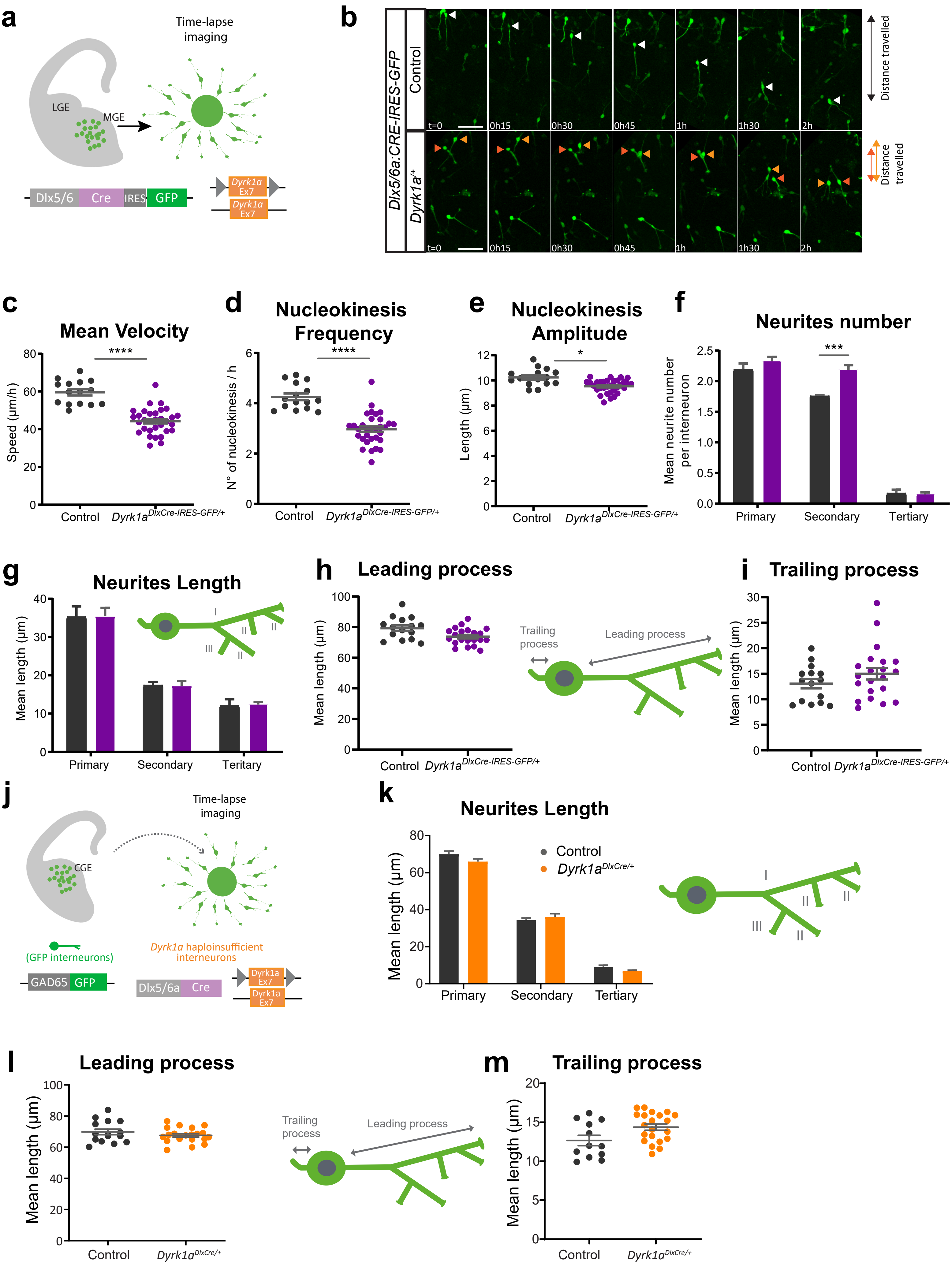

### Supplementary Fig.5

#### Supplementary Fig. 5

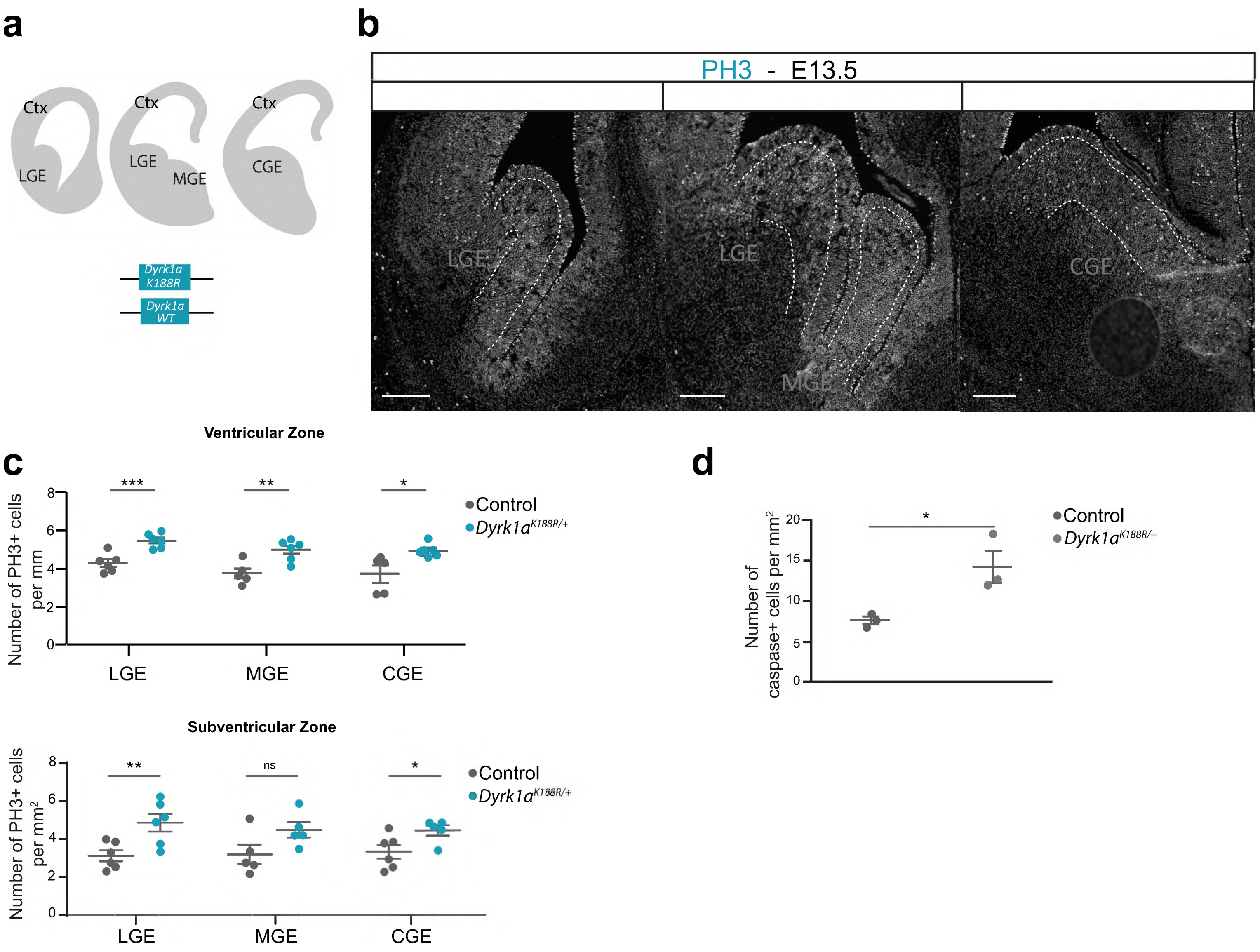

### Supplementary fig.7

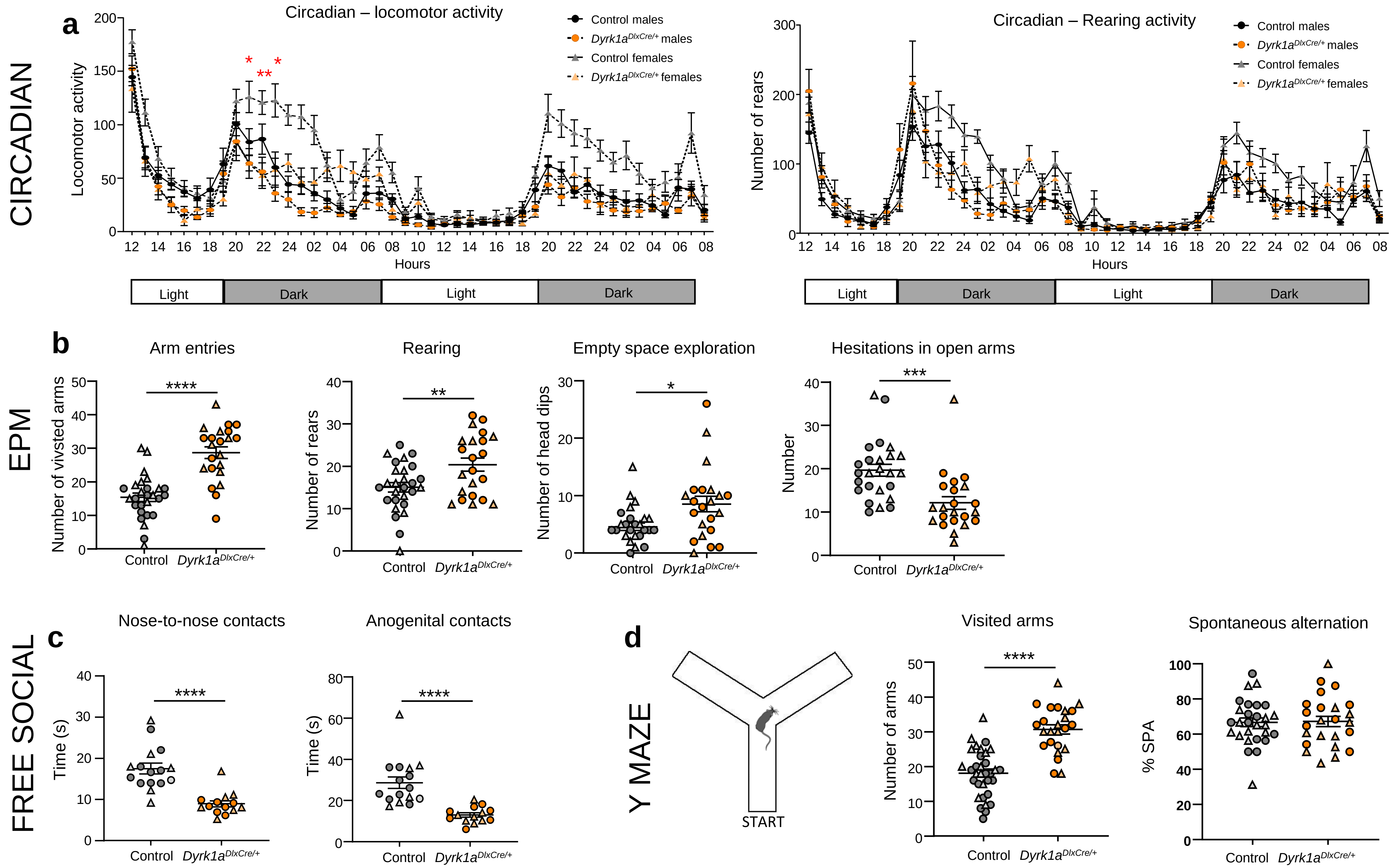
