## Supplementary fig.6 for "Interneuron migration defects during corticogenesis contribute to *Dyrk1a* haploinsufficiency syndrome pathogenesis via actomyosin dynamics deregulations"

a

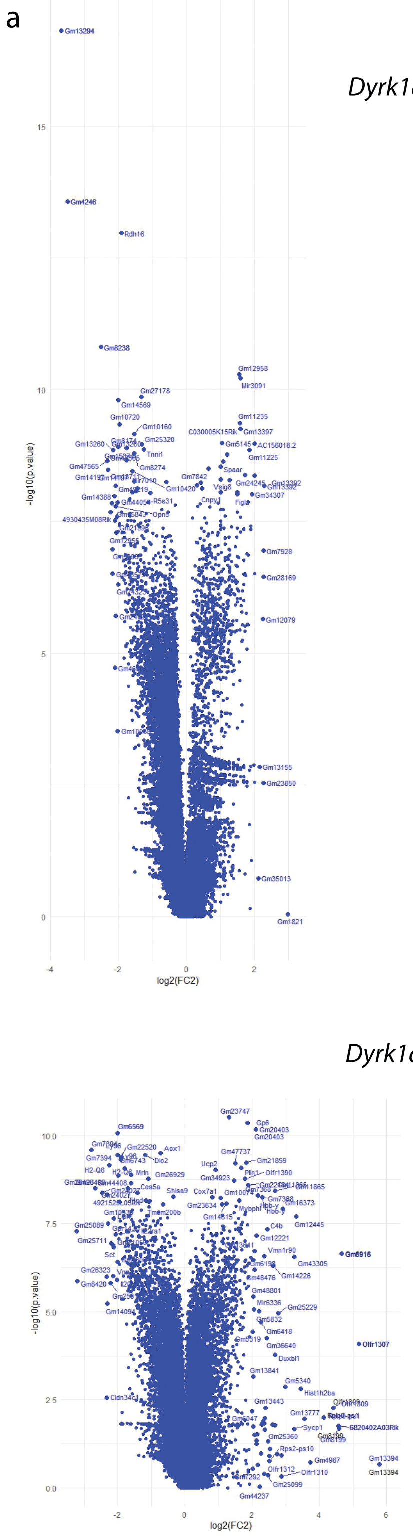


b

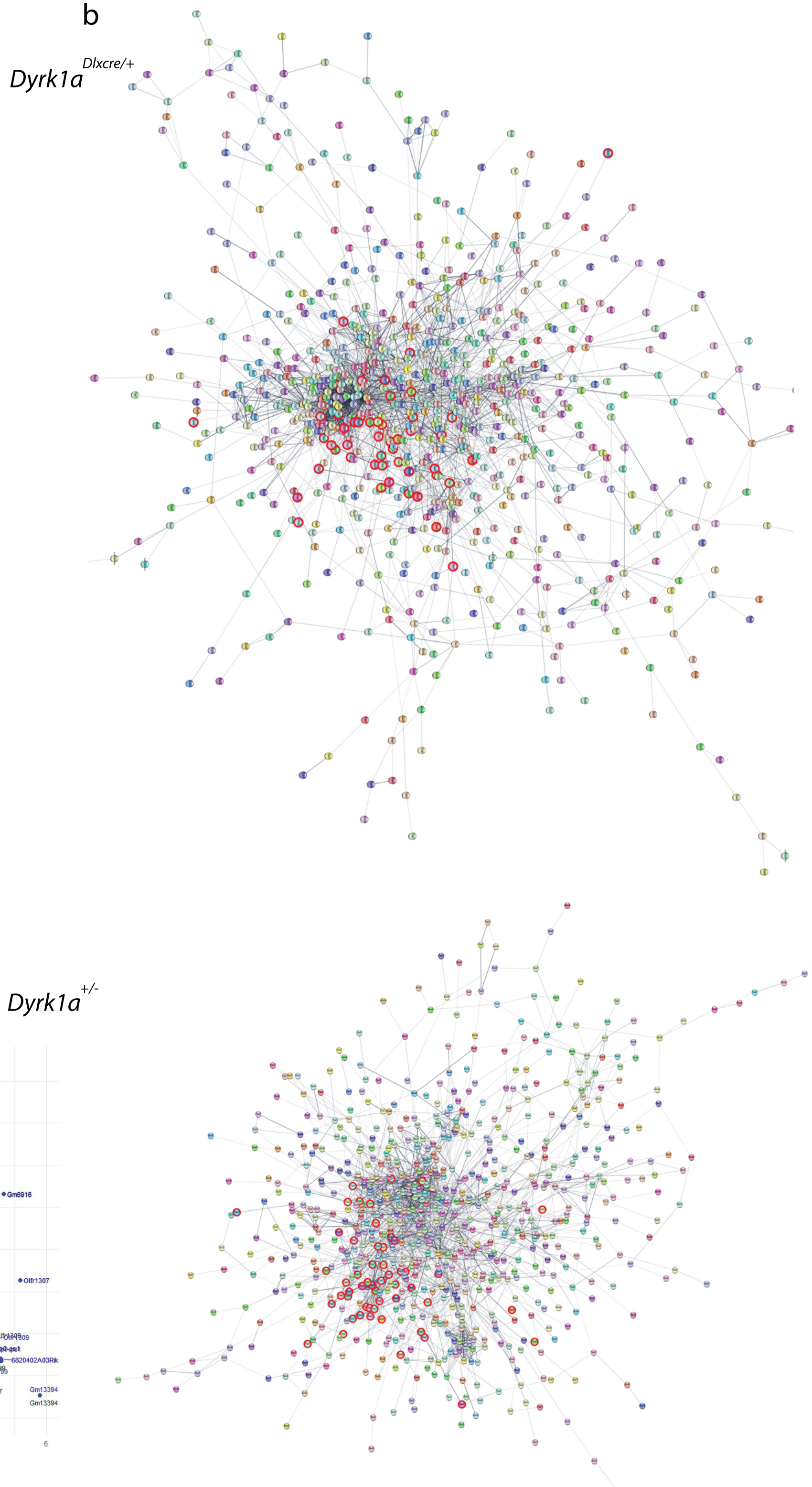


c

| *Dyrk1a* Genotype | # background genes | # genes | category | description | FDR value | genes |
|---|---|---|---|---|---|---|
| *Dlx/+* | 315 | 43 | Reactome Pathways | Class A/1 (Rhodopsin-like receptors) | 1.6E-4 | Drd3\|Cxcl13\|Lhcgr\|C3\|Gpr39\|F2\|Oxt\|Nmu\|Ppbp\|Prok2\|Rho\|Avp\|Trhr\|P2ry2\|Drd5\|Npffr2\|Hrh4\|Npbwr1\|C3ar1\|Cxcr3\|Ffar2\|Mtnr1b\|Oxgr1\|P2ry4\|Rxfp4\|Galr3\|Gphb5\|Htr1f\|Opn5\|Ptafr\|Gpr65\|Adrb3\|Htr1d\|Mc1r\|Npb\|Insl5\|Ccr5\|Ccrl2\|Hcrtr1\|Ccr9\|C5ar1\|Pyy\|C5ar2 |
| *Dlx/+* | 404 | 50 | Reactome Pathways | GPCR ligand binding | 1.6E-4 | Gprc6a\|Drd3\|Cxcl13\|Lhcgr\|C3\|Gpr39\|F2\|Oxt\|Nmu\|Ppbp\|Prok2\|Rho\|Avp\|Trhr\|P2ry2\|Drd5\|Npffr2\|Hrh4\|Npbwr1\|C3ar1\|Cxcr3\|Ffar2\|Mtnr1b\|Oxgr1\|P2ry4\|Rxfp4\|Galr3\|Gphb5\|Htr1f\|Opn5\|Tas2r137\|Casr\|Ptafr\|Gpr65\|Adrb3\|Htr1d\|Ramp1\|Mc1r\|Gcg\|Npb\|Insl5\|Ccr5\|Ccrl2\|Hcrtr1\|Ccr9\|C5ar1\|Grm6\|Pyy\|Crhbp\|C5ar2 |
| *Dlx/+* | 266 | 36 | WikiPathways | GPCRs, non-odorant | 2.5E-4 | Tmigd3\|Gprc6a\|Drd3\|Lhcgr\|Ccr1\|Trhr\|P2ry2\|Drd5\|Npffr2\|Npy6r\|Hrh4\|Npbwr1\|C3ar1\|Cxcr3\|Ffar2\|Mtnr1b\|Oxgr1\|P2ry4\|Rxfp4\|Gprc5a\|Htr1f\|Opn5\|Tas2r137\|Casr\|Ptafr\|Gpr65\|Adrb3\|Htr1d\|Mc1r\|Ccrl2\|Hcrtr1\|Ccr9\|C5ar1\|Grm6\|Ccr2\|C5ar2 |
| *+/-* | 315 | 43 | Reactome Pathways | Class A/1 (Rhodopsin-like receptors) | 1.6E-4 | Drd3\|Cxcl13\|Lhcgr\|C3\|Gpr39\|F2\|Oxt\|Nmu\|Ppbp\|Prok2\|Rho\|Avp\|Trhr\|P2ry2\|Drd5\|Npffr2\|Hrh4\|Npbwr1\|C3ar1\|Cxcr3\|Ffar2\|Mtnr1b\|Oxgr1\|P2ry4\|Rxfp4\|Galr3\|Gphb5\|Htr1f\|Opn5\|Ptafr\|Gpr65\|Adrb3\|Htr1d\|Mc1r\|Npb\|Insl5\|Ccr5\|Ccrl2\|Hcrtr1\|Ccr9\|C5ar1\|Pyy\|C5ar2 |
| *+/-* | 404 | 50 | Reactome Pathways | GPCR ligand binding | 1.6E-4 | Gprc6a\|Drd3\|Cxcl13\|Lhcgr\|C3\|Gpr39\|F2\|Oxt\|Nmu\|Ppbp\|Prok2\|Rho\|Avp\|Trhr\|P2ry2\|Drd5\|Npffr2\|Hrh4\|Npbwr1\|C3ar1\|Cxcr3\|Ffar2\|Mtnr1b\|Oxgr1\|P2ry4\|Rxfp4\|Galr3\|Gphb5\|Htr1f\|Opn5\|Tas2r137\|Casr\|Ptafr\|Gpr65\|Adrb3\|Htr1d\|Ramp1\|Mc1r\|Gcg\|Npb\|Insl5\|Ccr5\|Ccrl2\|Hcrtr1\|Ccr9\|C5ar1\|Grm6\|Pyy\|Crhbp\|C5ar2 |
| *+/-* | 266 | 36 | WikiPathways | GPCRs, non-odorant | 2.5E-4 | Tmigd3\|Gprc6a\|Drd3\|Lhcgr\|Ccr1\|Trhr\|P2ry2\|Drd5\|Npffr2\|Npy6r\|Hrh4\|Npbwr1\|C3ar1\|Cxcr3\|Ffar2\|Mtnr1b\|Oxgr1\|P2ry4\|Rxfp4\|Gprc5a\|Htr1f\|Opn5\|Tas2r137\|Casr\|Ptafr\|Gpr65\|Adrb3\|Htr1d\|Mc1r\|Ccrl2\|Hcrtr1\|Ccr9\|C5ar1\|Grm6\|Ccr2\|C5ar2 |

d

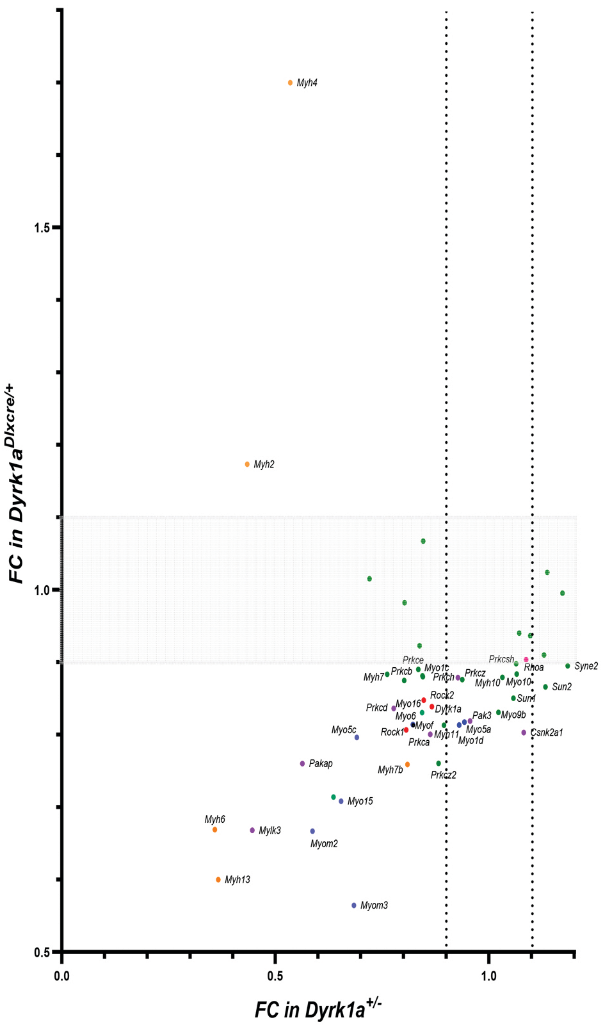
