## Supplementary material for "Interneuron migration defects during corticogenesis contribute to *Dyrk1a* haploinsufficiency syndrome pathogenesis via actomyosin dynamics deregulations": Table S1

|  | | *Dyrk1a^+/-^* | | | *Dyrk1a^Dlx/+^* | | |
| --- | --- | --- | --- | --- | --- | --- | --- |
| Ensembl ID | Gene name | FC | Pvalue | FDR | FC | Pvalue | FDR |
| ENSMUSG00000022897 | *Dyrk1a* | 0.87 | 1.69E-04 | 1.51E-03 | 0.84 | 6.11E-03 | 3.00E-02 |
| ENSMUSG00000020580 | *Rock2* | 0.85 | 2.04E-02 | 3.70E-02 | 0.85 | 2.04E-02 | 3.70E-02 |
| ENSMUSG00000024290 | *Rock1* | 0.81 | 1.63E-02 | 3.05E-02 | 0.81 | 1.63E-02 | 3.05E-02 |
| ENSMUSG00000031698 | *Mylk3* | 0.45 | 6.99E-09 | 5.53E-06 | 0.67 | 3.38E-02 | 5.81E-02 |
| ENSMUSG00000031284 | *Pak3* | 0.96 | 1.67E-02 | 4.48E-02 | 0.82 | 2.53E-02 | 4.48E-02 |
| ENSMUSG00000021108 | *Prkch* | 0.93 | 3.42E-02 | 7.86E-02 | 0.88 | 4.33E-02 | 7.27E-02 |
| ENSMUSG00000021948 | *Prkcd* | 0.78 | 6.18E-05 | 7.62E-04 | 0.84 | 9.99E-03 | 3.00E-02 |
| ENSMUSG00000050965 | *Prkca* | 0.86 | 6.31E-05 | 7.72E-04 | 0.80 | 8.74E-03 | 3.00E-02 |
| ENSMUSG00000074698 | *Csnk2a1* | 1.08 | 1.54E-03 | 7.34E-03 | 0.80 | 1.52E-03 | 1.32E-02 |
| ENSMUSG00000007815 | *Rhoa* | 1.09 | 1.84E-03 | 8.39E-03 | 0.90 | 5.32E-01 | 6.13E-01 |
| ENSMUSG00000029516 | *Cit* | 1.10 | 2.34E-02 | 5.81E-02 | 0.94 | 7.91E-01 | 8.34E-01 |
| ENSMUSG00000030602 | *Pak4* | 1.14 | 4.36E-04 | 2.92E-03 | 1.02 | 1.62E-01 | 2.36E-01 |
| ENSMUSG00000038683 | *Pak1ip1* | 1.13 | 1.84E-03 | 8.38E-03 | 0.91 | 8.71E-01 | 9.01E-01 |
| ENSMUSG00000074923 | *Pak6* | 0.84 | 5.20E-04 | 3.31E-03 | 0.92 | 6.70E-01 | 7.38E-01 |
| ENSMUSG00000089945 | *Pakap* | 0.56 | 2.58E-04 | 2.04E-03 | 0.76 | 2.86E-01 | 3.81E-01 |
| ENSMUSG00000003402 | *Prkcsh* | 1.06 | 5.17E-03 | 1.81E-02 | 0.90 | 5.44E-01 | 6.24E-01 |
| ENSMUSG00000026778 | *Prkcq* | 0.72 | 2.49E-05 | 4.18E-04 | 1.02 | 1.11E-01 | 1.70E-01 |
| ENSMUSG00000029053 | *Prkcz* | 0.94 | 4.09E-02 | 9.03E-02 | 0.88 | 5.65E-02 | 9.19E-02 |
| ENSMUSG00000045038 | *Prkce* | 0.83 | 3.57E-06 | 1.30E-04 | 0.89 | 1.40E-01 | 2.06E-01 |
| ENSMUSG00000052889 | *Prkcb* | 0.80 | 1.53E-05 | 3.02E-04 | 0.87 | 5.10E-01 | 5.93E-01 |
| ENSMUSG00000078816 | *Prkcg* | 0.84 | 1.76E-02 | 4.66E-02 | 0.88 | 5.88E-02 | 9.50E-02 |
| ENSMUSG00000108314 | *Prkcz2* | 0.88 | 3.54E-03 | 1.36E-02 | 0.76 | 5.11E-02 | 8.42E-02 |
| ENSMUSG00000024387 | *Csnk2b* | 1.07 | 2.55E-02 | 6.23E-02 | 0.94 | 3.11E-01 | 4.08E-01 |
| ENSMUSG00000031461 | *Myom2* | 0.59 | 6.42E-04 | 3.86E-03 | 0.67 | 2.27E-03 | 1.78E-02 |
| ENSMUSG00000033577 | *Myo6* | 0.82 | 1.49E-03 | 7.17E-03 | 0.81 | 2.56E-02 | 4.53E-02 |
| ENSMUSG00000033590 | *Myo5c* | 0.69 | 1.53E-03 | 7.30E-03 | 0.80 | 1.74E-02 | 3.23E-02 |
| ENSMUSG00000034593 | *Myo5a* | 0.94 | 8.07E-03 | 2.57E-02 | 0.82 | 6.56E-04 | 7.26E-03 |
| ENSMUSG00000035441 | *Myo1d* | 0.93 | 7.30E-03 | 2.37E-02 | 0.81 | 7.52E-03 | 3.00E-02 |
| ENSMUSG00000037139 | *Myom3* | 0.68 | 2.13E-02 | 5.41E-02 | 0.56 | 2.56E-06 | 1.47E-04 |
| ENSMUSG00000042678 | *Myo15* | 0.65 | 5.43E-05 | 6.99E-04 | 0.71 | 1.48E-02 | 3.00E-02 |
| ENSMUSG00000048612 | *Myof* | 0.82 | 1.77E-02 | 4.69E-02 | 0.81 | 3.69E-03 | 2.46E-02 |
| ENSMUSG00000004677 | *Myo9b* | 1.02 | 1.27E-02 | 3.65E-02 | 0.83 | 1.96E-01 | 2.78E-01 |
| ENSMUSG00000017774 | *Myo1c* | 0.85 | 2.97E-03 | 1.20E-02 | 0.88 | 5.65E-01 | 6.43E-01 |
| ENSMUSG00000022272 | *Myo10* | 1.06 | 9.55E-03 | 2.92E-02 | 0.88 | 8.19E-01 | 8.56E-01 |
| ENSMUSG00000025401 | *Myo1a* | 1.17 | 4.13E-03 | 1.53E-02 | 1.00 | 6.98E-01 | 7.62E-01 |
| ENSMUSG00000039057 | *Myo16* | 0.84 | 3.07E-05 | 4.78E-04 | 0.83 | 6.64E-02 | 1.06E-01 |
| ENSMUSG00000042064 | *Myo3b* | 0.80 | 3.82E-04 | 2.66E-03 | 0.98 | 9.52E-01 | 9.64E-01 |
| ENSMUSG00000072720 | *Myo18b* | 0.85 | 2.99E-04 | 2.26E-03 | 1.07 | 1.23E-01 | 1.86E-01 |
| ENSMUSG00000033196 | *Myh2* | 0.43 | 3.27E-06 | 1.23E-04 | 1.17 | 5.59E-03 | 3.00E-02 |
| ENSMUSG00000040752 | *Myh6* | 0.36 | 4.21E-08 | 1.30E-05 | 0.67 | 4.85E-04 | 5.72E-03 |
| ENSMUSG00000057003 | *Myh4* | 0.53 | 6.20E-05 | 7.64E-04 | 1.70 | 1.63E-02 | 3.05E-02 |
| ENSMUSG00000060180 | *Myh13* | 0.37 | 2.22E-07 | 2.70E-05 | 0.60 | 7.00E-03 | 3.00E-02 |
| ENSMUSG00000074652 | *Myh7b* | 0.81 | 6.31E-03 | 2.12E-02 | 0.76 | 4.27E-03 | 2.73E-02 |
| ENSMUSG00000018830 | *Myh11* | 0.90 | 4.68E-03 | 1.68E-02 | 0.81 | 2.65E-01 | 3.59E-01 |
| ENSMUSG00000020900 | *Myh10* | 1.03 | 7.07E-03 | 2.31E-02 | 0.88 | 3.24E-01 | 4.23E-01 |
| ENSMUSG00000053093 | *Myh7* | 0.76 | 8.62E-03 | 2.70E-02 | 0.88 | 1.96E-01 | 2.78E-01 |
| ENSMUSG00000055775 | *Myh8* | 0.64 | 1.02E-02 | 3.07E-02 | 0.71 | 9.12E-02 | 1.41E-01 |
| ENSMUSG00000030672 | *Mylpf* | 1.47 | 5.33E-04 | 3.37E-03 | 1.09 | 2.34E-01 | 3.23E-01 |
| ENSMUSG00000061816 | *Myl1* | ND |  |  | 0.62 | 1.17E-02 | 3.00E-02 |
| ENSMUSG00000113178 | *Mylf-ps* | ND |  |  | 0.60 | 2.85E-03 | 2.04E-02 |
| ENSMUSG00000036817 | *Sun1* | 1.06 | 6.70E-04 | 3.98E-03 | 0.85 | 3.54E-01 | 4.53E-01 |
| ENSMUSG00000042524 | *Sun2* | 1.13 | 5.52E-05 | 7.07E-04 | 0.87 | 3.61E-01 | 4.61E-01 |
| ENSMUSG00000063450 | *Syne2* | 1.18 | 9.91E-06 | 2.34E-04 | 0.89 | 9.15E-01 | 9.36E-01 |

Table S3: Selected genes involved in nucleokinesis and cytoskeleton reorganisation found misregulated in DYRK1A mutants (ND not detected; based on (Javier-Torrent and Saura, 2020; Blake et al., 2021)
